# Continuous optical detection of small-molecule analytes in complex biomatrices

**DOI:** 10.1101/2023.03.03.531030

**Authors:** Amani A. Hariri, Alyssa P. Cartwright, Constantin Dory, Yasser Gidi, Steven Yee, Kaiyu Fu, Kiyoul Yang, Diana Wu, Ian A. P. Thompson, Nicolò Maganzini, Trevor Feagin, Brian E. Young, Behrad Habib Afshar, Michael Eisenstein, Michel Digonnet, Jelena Vuckovic, H. Tom Soh

## Abstract

Current technology for measuring specific biomarkers – continuously in complex samples, without sample preparation – is limited to just handful of molecules such as glucose and blood oxygen. In this work, we present the first optical biosensor system that enables continuous detection of a wide range of biomarkers in complex samples, such as human plasma. Our system employs a modular duplex-bubble switch (DBS) architecture that converts aptamers into structure-switching fluorescence probes whose affinity and kinetics can be readily tuned. These DBS constructs are coupled to a fiber-optic detector that measures the fluorescence change only within an evanescent field, thereby minimizing the impact of background autofluorescence and enabling direct detection of analytes at physiologically relevant concentrations even in interferent-rich sample matrices. Using our system, we achieved continuous detection of dopamine in artificial cerebrospinal fluid for >24 hours with sub-second resolution and a limit of detection (LOD) of 1 µM. We subsequently demonstrated the system’s generalizability by configuring it to detect cortisol with nanomolar sensitivity in undiluted human plasma. Both sensors achieved LODs orders of magnitude lower than the *K*_*D*_ of the DBS element, highlighting the potential to achieve sensitive detection even when using aptamers with modest affinity.

## Introduction

The ability to achieve continuous detection of specific biomarkers in complex samples offers exciting potential for both biological research and clinical medicine. The advent of continuous glucose monitoring (CGM) technologies offers one current example, giving patients a powerful tool for the management of type 1 diabetes, with well-validated benefits in terms of clinical outcomes.^1^ Unfortunately, such capabilities are currently only available for a small number of analytes, because existing technologies employ detection modalities that are not generalizable to other biomarkers. For example, CGM systems rely on a glucose-specific enzymatic reaction to generate a readout.^2^ Recent years have seen considerable effort to develop generalizable sensor architectures that can be utilized to detect a wide range of analytes, including the use of electrochemical sensors based on structure-switching aptamers.^3,4^ The first such “real-time” sensor was described in 2009 by Swensen *et al*., who demonstrated the continuous detection of micromolar concentrations of cocaine in undiluted serum with minute-scale temporal resolution.^5^ In subsequent work, Ferguson and colleagues described an improved version of this electrochemical aptamer sensor design, which enabled the continuous detection of multiple small-molecule drugs in the circulation of live rats with sub-minute temporal resolution over the course of several hours.^6^ More recently, a number of other research groups have published additional innovative demonstrations of the use of electrochemical aptamer-based sensors for continuous detection^7,8^ These sensors employ engineered aptamers that undergo a conformation change upon target binding. These are labeled at one end with an electrochemical reporter and then immobilized onto an electrode at the other terminus, such that a change in target concentration produces a proportional change in the redox current by altering the interaction between the redox tag and the electrode surface. These sensors can be integrated with standard electronic components, enabling rapid and simple signal analysis and interpretation.^9^

However, stable and sensitive analyte detection in complex samples over extended periods of time generally remains a major challenge with electrochemical sensors, as outlined in previous work.^10^ First, since detection is directly predicated on the ability of the redox reporter to interact with the electrode surface, electrode-based sensors are vulnerable to fouling from interferents present in the sample. Surface passivation strategies can mitigate this problem to some extent,^11^ but fouling will inevitably occur when these sensors are deployed in complex mixtures over extended periods of time. Additionally, the electrochemical current change produced by individual target binding events can be relatively small and difficult to resolve for analytes that are present at low physiological concentrations.

As an alternative strategy that can potentially overcome these challenges, we have developed a platform for sensitive, continuous molecular detection in complex samples that couples structure-switching aptamer probes with an optical rather than an electrochemical read-out. Optical detection offers the advantage that the signaling mechanism need not be dependent on direct interaction of either the analyte or the aptamer with the sensor surface, and thus should be less susceptible to the effects of surface fouling. For example, one can exploit proximity-based intermolecular phenomena such as fluorophore-quencher interactions or Förster resonance energy transfer (FRET). In addition, photonic detection generally offers higher sensitivity than electronic detection due to the commercial availability of devices such as single-photon detectors, offering the potential to achieve higher sensitivity using the same aptamer probes.

Our system employs two key technologies: a novel duplex-bubble switch (DBS) aptamer switch architecture that is designed for continuous optical reporting, and a fiber-optic probe integrated with single-photon-counting hardware to achieve highly sensitive detection. This probe design exploits an evanescent field-based sensing approach that enables direct measurement of analytes in complex matrices such as plasma by rejecting background autofluorescence from interferents in the sample, while also minimizing the impact of fouling. We first show mathematically and experimentally that we can fine-tune the DBS design to precisely adjust the thermodynamics and kinetics of the resulting aptamer switches. We subsequently generated a DBS-based dopamine sensor that achieved robust continuous detection of dopamine in both buffer and artificial cerebrospinal fluid (aCSF) for more than a day with fast time resolution (less than 3 s) and a dynamic range spanning 500 nM–500 μM. To demonstrate the generalizability of our approach, we subsequently developed a cortisol-specific DBS probe and integrated it with our optical detection hardware. This sensor achieved cortisol detection with nanomolar sensitivity and a dynamic range suitable for detecting physiologically relevant concentrations (200 nM–100 μM) for multiple hours in undiluted human plasma.

## Results and Discussion

### Duplex bubble switch (DBS) design and modeling

The DBS is a double-stranded, multi-domain fluorescent sensor (**Fig. 1**), in which the aptamer is hybridized to a complementary displacement strand sequence. The aptamer is modified with an internal fluorophore, which is positioned to be adjacent to a quencher tag coupled to the 5’ terminus of the displacement strand. The 5’ end of the aptamer and the 3’ end of the displacement strand are each connected to complementary ‘anchor domains’ via unpaired sequences that form the ‘bubble domain’. This latter element acts as a flexible spacer to prevent conformational changes that would hinder aptamer target recognition and binding, while the anchor domain serves to tether the DBS to the sensor substrate.

**Figure 1:**
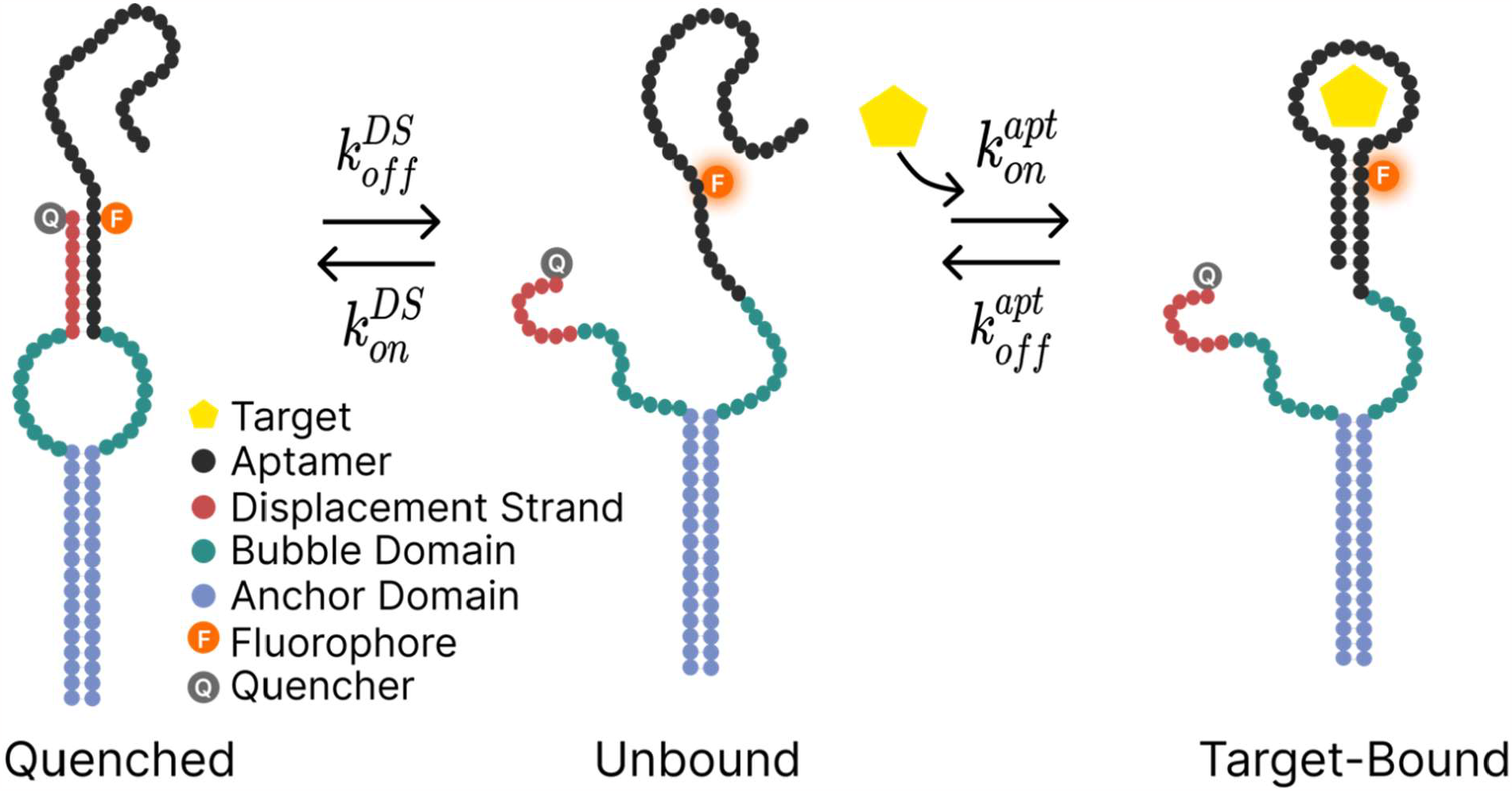
Schematic of the duplex bubble switch (DBS). The DBS converts an existing aptamer into a switch that produces a target-concentration-dependent signal based on alterations in the interaction of a fluorophore-quencher pair. In the absence of the aptamer target, the DBS exists in an equilibrium state between its quenched form (left), in which the quencher and fluorophore are in close proximity due to binding of the displacement strand to the aptamer, and the unbound state (middle), in which the two strands are dissociated. In the presence of the target, the system shifts to an equilibrium between the unbound state and the target-bound state (right), in which the fluorophore signal is greatly enhanced. This results in a target concentration-dependent signal increase.

The DBS can exist in three states: a quenched state in which the displacement strand is bound to the aptamer and thus suppresses the fluorophore signal, an unfolded state, and a target-bound state. In the absence of target, the quenched and unfolded states are in equilibrium, defined by the constant *K*_Q_. Target binding shifts the equilibrium to the right, favoring the formation of the target-bound state, in which the quencher and fluorophore are now physically separated, over the quenched state, thereby generating a signal that is proportional to the target concentration. The DBS design offers the advantage of enabling simple, rational design without requiring prior knowledge of the parent aptamer’s secondary structure or binding site, thereby offering a more generalizable approach to the design of tunable aptamer switches. Indeed, any aptamer generated via Capture-SELEX^12-14^ can be readily incorporated into a DBS construct, because the 5’ end of the aptamer domain is hybridized to its displacement strand via the same complementary sequence used during the original Capture-SELEX experiment, with no involvement from the target-binding domain of the aptamer.

We can also tune DBS thermodynamics and kinetics by modulating the hybridization of the aptamer to the displacement strand, enabling the sensor to achieve a dynamic range that is optimal for the desired target concentration while also undergoing switching at physiologically-relevant concentrations.^15,16^ The DBS design offers two parallel mechanisms for sensor tuning: decreasing the length of the bubble domain (L_bubble_) is expected to reduce binding affinity (*i*.*e*., increase K_D_^eff^) while decreasing background and increasing the overall rate of binding, whereas decreasing the length of the displacement strand (L_DS_) reduces K_D_^eff^ while increasing background signal and increasing the rate of binding. We employed a mathematical model to assess the degree of tunability that could be achieved by manipulating these two parameters. Example calculations for this model can be found in Reference 16, and a full derivation can be found in **Supplementary Note 1**.

### Assessing the affinity and binding response of a dopamine DBS

As an initial demonstration for this work, we constructed a DBS based on an existing dopamine aptamer^12^ (39 nt, *K*_*D*_ = 150 nM) and tested these thermodynamic and kinetic tuning principles with our dopamine DBS in solution (**Supplementary Table 1**). We generated an array of DBS switches incorporating displacement strands with L_DS_ ranging from 7 to 12 nucleotides (nt) and L_bubble_ ranging from 12 to 32 nt, and tested the dopamine-binding affinity of these constructs using a plate reader-based assay (see Methods). We describe our switches using the nomenclature of L_bubble_-L_DS_, such that a construct with a 22-nt bubble and a 7-bp displacement strand is referred to as 22-7. We selected a Cy3 fluorophore and an Iowa Black quencher to label the aptamer and displacement strands, respectively, in order to achieve long-term measurement with minimal fluorescence background or photobleaching. As expected, increasing L_DS_ on its own resulted in a lower background signal and lower apparent dopamine affinity. For instance, as we increased L_DS_ from 7 to 12 nt while maintaining L_bubble_ = 22, the majority of the corresponding aptamer switches exhibited lower sensitivity, with an increase in K_Deff_ from 35 μM to 170 μM (**Fig. 2a**). In contrast, we observed greater DBS sensitivity as the size of L_bubble_ increased from 12 to 22 or 32 nt, with significant improvement in K_D_^eff^ for every 10 additional bases inserted (K_D_^eff^ = 177.1, 68.31 and 48.89 μM for L_bubble_ = 12, 22, and 32 nt, respectively) while maintaining a constant L_DS_ = 10 (**Supplementary Figure S1, S2**; **Supplementary Table 2)**.

**Figure 2:**
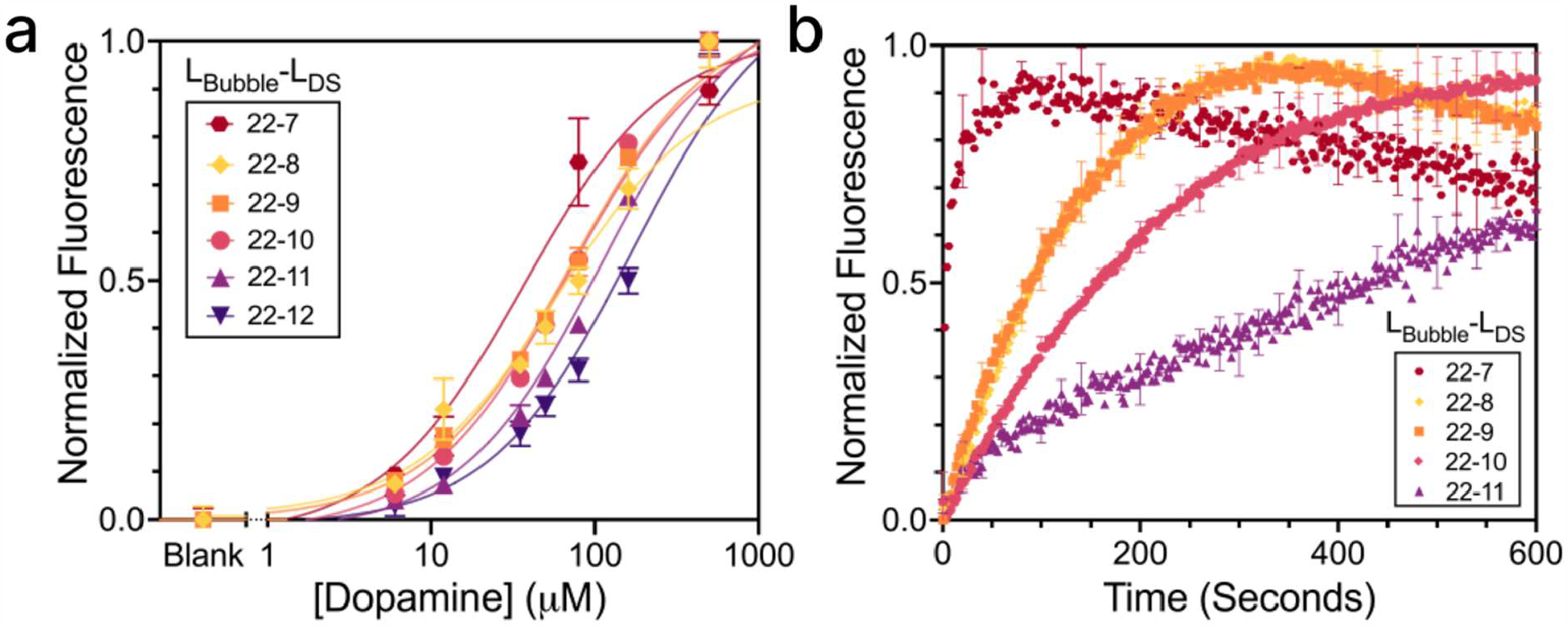
Tuning the performance of a dopamine DBS in solution. **a)** Changing the LDS of our DBS from 7 to 12 bp while maintaining L_bubble_ at 22 nt shifts the binding curve to the right and reduces background signal. **b**) Modulating temporal response to an injection of 50 µM dopamine via tuning of L_DS_. Increasing L_DS_ from 7 to 11 bp with a fixed L_bubble_ of 22 nt results in slower kinetics. Error bars are presented for every 10^th^ data point for better visualization. Error bars in **a** and **b** represent the standard deviation of the average (n = 3). Raw thermodynamic plots for all DBS constructs are provided in **Supplementary Figures S1** and **S2**. Data normalization and fits are explained in Methods.

We subsequently tested our ability to tune the temporal response of our various constructs by adding 50 µM dopamine to plate wells containing 250 nM of each DBS construct and then observing the kinetics of the fluorescent response. Decreasing L_bubble_ with a constant L_DS_ did not meaningfully alter the kinetic response, but decreasing L_DS_ with a constant L_bubble_ (**Supplementary Figure S3, Supplementary Table 3**) resulted in significantly faster responses. We could dramatically increase temporal resolution (the switching time constant *τ*_*obs*_ = 1/*K*_*obs*_) by over 300-fold—from ∼17 min to less than 3 s— as we decreased L_DS_ from 11 to 7 nt in constructs with L_bubble_ = 22 (**Fig. 2b)**. For constructs with longer L_DS_, we hypothesized that it should be possible to further improve their kinetic response by introducing mismatches into the aptamer-displacement strand duplex. Previous work has shown that such mismatches can enable more precise control over the binding curve and enhance the ability to increase kinetics without affecting affinity.^16,17^ We therefore introduced single-base mismatches at different positions within the hairpin of the parent aptamer sequence and tested it with multiple L_DS_ constructs as L_DS_ = 11 nt, a design that displayed very slow kinetics in the absence of mismatches (**Supplementary Figure S3**). In most cases, the introduction of mismatches dramatically increased the DBS association rate constant (k_on_). For example, for construct 12-11, a mismatch at position 8 in the aptamer shifted k_on_ from ∼8 min to less than 35 s—a >12-fold improvement.

Since the DBS constructs will be coupled to a solid support in our biosensor system, we next assessed the effect of surface anchoring on the affinity and binding kinetics of our 22-7 construct. We determined that 22-7 was the best candidate for further development, as it achieved the highest affinity of any of the DBS designs tested (K_D_ = 35 µM) with a good dynamic range (1– 100 µM) while also undergoing switching at fast timescales (seconds). We biotinylated the 3’ end of the displacement strand-coupled anchor strand for surface attachment, and modified its 5’ end with Iowa Black quencher; the aptamer strand was internally modified with a Cy3 reporter. We immobilized this modified 22-7 construct onto a passivated, streptavidin-coated glass surface and used total internal reflection fluorescence (TIRF) microscopy to image dopamine binding with single-molecule resolution^18^ (**Fig. 3a**).

**Figure 3:**
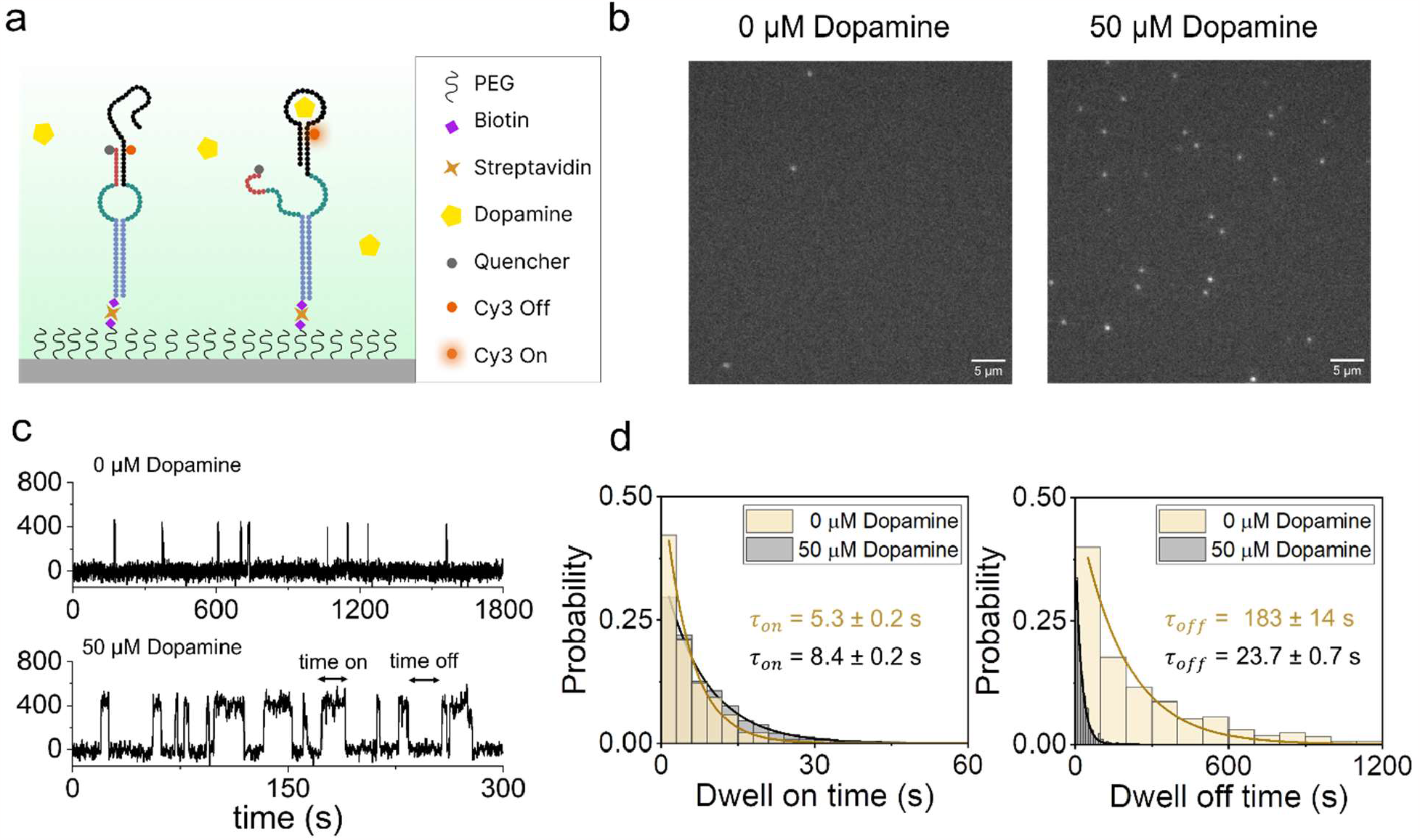
Characterization of surface-coupled dopamine DBS constructs. **a)** Schematic of DBS coupling to a streptavidin-modified glass substrate for analysis via total internal reflection fluorescence (TIRF) microscopy. **b)** The interaction of dopamine with surface-attached DBS probes yields temporal patterns of repeated binding and dissociation. Panels show single movie frames from a microscope field of view before (left) and after (right) dopamine addition, with bright puncta at locations where single fluorescent probes are bound. **c**) Representative intensity-time trajectories indicating binding/unbinding to a single DBS. **d**) Histograms of τ_on_ (left) and τ_off_ (right) for all intensity-versus-time trajectories observed within a single field of view in the presence or absence of dopamine with mono-exponential fits.

As we applied the dopamine-containing solution to the glass surface, we were able to visualize in real-time the binding and release of individual dopamine molecules from the DBS as sharp increases and decreases in intensity (**Fig. 3b, Supplementary Figure S4**). Our results confirmed that our DBS sensor can achieve robust dopamine detection when anchored to a solid substrate. Representative fluorescence intensity trajectories are shown in **Figure 3c** and **Supplementary Figure S4**. By monitoring individual binding events in real-time, we were able to calculate *k*_on_ via 1/τ_off_ = *k*_on_*c*, where *c* is the dopamine concentration and τ_off_ is the mean dwell off-time. Similarly, the dissociation rate constant (*k*_off_) can be calculated via 1/τ_on_ = *k*_off_, where τ_on_ is the mean dwell on-time. In the absence of target, we observed mono-exponential decays indicating long dwell off-time (183 s) and sporadic and short on-times (5.3 s). In contrast, the addition of target yielded mono-exponential decays, with a marked decrease in dwell off-times (23.7 s) and an increase in dwell on-times (8.40 s) (**Fig. 3d**). These equilibria in the absence and presence of dopamine allowed us to estimate a K_D_^eff^ of 148 µM (K_D_^eff^ = τ_off_ /τ_on_), which is three-fold higher than the measurement we obtained in solution (K_D_^eff^ = 35 µM). These results indicated that surface coupling was influencing DBS affinity, possibly due to factors such as probe accessibility, the distance between probes, or probe-surface interactions^19^.

### Development of a near-field optical probe system

To measure the signals from the DBS probes, we developed a fiber probe-based optical detection system that can achieve sensitive analyte detection directly in complex biological samples such as plasma without any sample preparation. The design of our optical probe system was guided by three major goals: 1) rejection of autofluorescence from the sample, 2) prevention of non-specific binding and biofouling, and 3) maximizing sensitivity.

We designed our fiber probe to monitor DBS binding based on fluorophore excitation within an evanescent field, which means that out-of-focus fluorescence and autofluorescence from complex samples are mostly rejected (**Fig. 4a**). The evanescent field is generated when total internal reflection of light occurs within the core of the optical fiber, which is coated with a lower refractive index material. The field decays as it extends into the medium perpendicular to the fiber surface, such that fluorophores immediately adjacent to the surface (typically within 100–1,000 nm)^20^ will be excited by the evanescent wave; part of the resulting emitted fluorescence will in turn be coupled back into the fiber and can subsequently be measured. As such, DBS constructs coupled to the probe surface will generate measurable fluorescent signals in response to target binding that are proportional to the target concentration, whereas unbound molecules will contribute very little to the measured signal, resulting in greatly reduced background noise.

**Figure 4:**
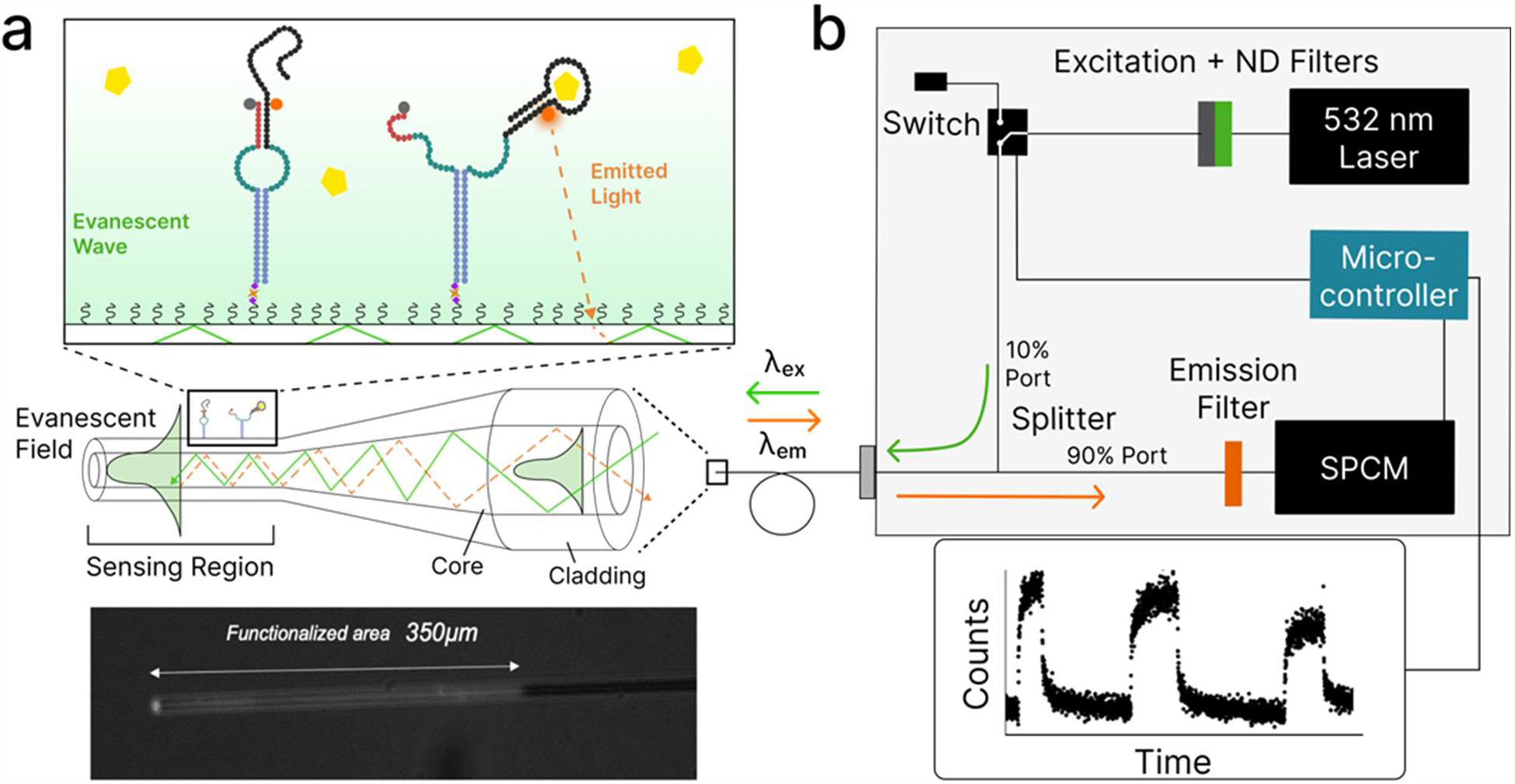
Near-field optical probe development and characterization. **a)** DBS constructs are immobilized onto a tapered optical fiber probe via biotin-avidin linkages to allow efficient near-field detection. Binding of target molecules to the DBS constructs is monitored in real time based on the excitation of fluorophore molecules within the evanescent field, where the emitted signal scales with incident power. Bottom panel shows fluorescence microscopy image of the tapered fiber tip functionalized with Cy3-labelled aptamers. **b)** A compact hardware system is used to excite DBS fluorescence and collect emitted fluorescence using a highly sensitive single-photon counting module (SPCM). Bottom panel shows an example of DBS signal over time as monitored in continuous mode.

In order to minimize biofouling, we functionalized the fiber tip in a multi-step process (see Methods). After cleaning with piranha, acetone, and water, the probes were functionalized with a mixture of unmodified and biotin-modified polyethylene glycol (PEG) polymer chains in order to insulate the probe against non-specific binding while also enhancing the specific attachment of DBS molecules to the surface via biotin/avidin interactions.^21^ We situated the probe within a microfluidic chamber (∼40 µL) by pressing a polycarbonate film with an adhesive gasket onto the PEG/PEG-biotin-coated fiber. Two silicone connectors were glued onto the predrilled holes of the film and served as inlet and outlet ports to manually dispense and wash out different solutions (**Supplementary Figure S5)**. The fibers were then incubated with a neutravidin solution, which enabled direct coupling of the biotinylated DBS molecules to the biotin moieties at the end of the PEG chains (**Fig. 4a**). Control experiments in which we incubated the functionalized probes with Cy3-labeled DNA strand in the absence of neutravidin ruled out any meaningful nonspecific binding of the aptamers to the sensor surface (**Supplementary Figure S6**), offering evidence that this design approach should generally help insulate against non-specific probe surface fouling.

Finally, to maximize our probe’s sensitivity, we optimized the fiber design for high transmission of fluorescent signal and efficient evanescent coupling to the DBS probes. We selected a commercially available, graded-index, multimode silica optical fiber with a core diameter of 62.5 µm, low attenuation (∼3-5 dB/km at wavelengths of 650-850 nm), and low background fluorescence in the emission band of our dyes (**Supplementary Figure S7**). The chemical robustness and lack of oxidation of silica-based materials means that they remain stable in aqueous media for prolonged periods of time, and silica-based surfaces can easily be modified with passivation treatments that minimize biofouling in complex media^22-25^. To maximize the sensing area—and therefore enhance the coupling between modes inside the fiber and the DBS construct—we tapered the base fiber down to a micrometer-scale tip (See Methods).^26^ Our fiber-tapering process was highly consistent, resulting in reproducible taper quality and dimensions and similar signals across several fiber probes (Signal CV= 0.10) (See **Supplementary Figure S8**).

This optical fiber probe was then coupled to a sensitive single-photon counting module (SPCM) to achieve optimal detection performance. The functionalized fiber was connected to an optoelectronic system (**Fig. 4b**) that is designed to both excite the dye molecules on the DBS and collect the resulting fluorescent signal. The excitation-detection module contains the excitation laser (532 nm), with the appropriate laser line filters including variable neutral density (ND) filters to control incident power to the fiber probe. The laser light is coupled into a fiber and fiber switch, which allows for interval measurements and is controlled via a microcontroller (Arduino). This enables the laser to be switched between the sample fiber path (ON) and a dead-end port (OFF). The excitation laser is coupled to the functionalized fiber through the 10% port of a 90:10 fiber coupler. The emission light of the biosensor is collected with the same fiber and is coupled back into the excitation/detection box through the 90% port and then spectrally separated from the excitation laser via long-pass and notch filters. The emission signal from the fiber probe is then directed to the SPCM, which is interfaced to the Arduino to record detection events and monitor fluorescence intensity over time. Data acquisition is performed by a LabVIEW GUI with an integration time of 500 ms and 500 nW output laser power to minimize photobleaching (**Supplementary Figure S9**). The fluorescence response of the aptamer switch is defined as the fluorescence intensity of the target after subtracting the background from a blank fiber. By correlating this signal to analyte concentration, we can achieve precise real-time measurements of analyte concentration changes over time.

### Continuous detection of dopamine in buffer and artificial cerebrospinal fluid

To characterize the sensitivity of the platform, we exposed our sensor to increasing and decreasing dopamine concentrations in buffer by intermittently injecting samples containing different analyte concentrations (0, 1, 8, 80, 200, or 800 μM) into the chamber. We observed a clear and proportional signal gain (**Fig. 5a, Supplementary Figure S10**), ranging from 29% at 1 μM to ∼2,000% at 800 μM, with an average CV of 0.15 at each concentration. We fitted the probe’s signal gain to a Langmuir isotherm and obtained a *K*_*D*_ of 231 μM, with a measured dynamic range of 1 μM to 800 μM **(Fig. 5a, Supplementary Table 4)**. This limit of detection (LOD) is more than two orders of magnitude below the *K*_*D*_ of the DBS itself (231 μM), demonstrating that even a DBS with modest affinity can achieve sensitive analyte detection in this probe design. By increasing the incident laser power—and thereby improving the signal-to-noise ratio—we could further lower the LOD from 1 μM to ∼200 nM (**Supplementary Figure S11**). However, the gains from this strategy are mitigated by fluorophore saturation and photobleaching, which would limit the duration of measurements that can be collected in this fashion (**Supplementary Figure S9**). We next characterized the kinetics of the sensor upon exposing the probe to 8, 80, 200, and 800 μM dopamine, and observed a rapid kinetic response, with k_on_ = 138 M^−1^s^−1^ and k_off_ = 0.102s^−1^. These measurements indicate that the probe reached 50% of its maximum signal within 7 s and then returned to 50% of baseline within 5.5 s (**Supplementary Figure S12, Supplementary Table 5**).

**Figure 5:**
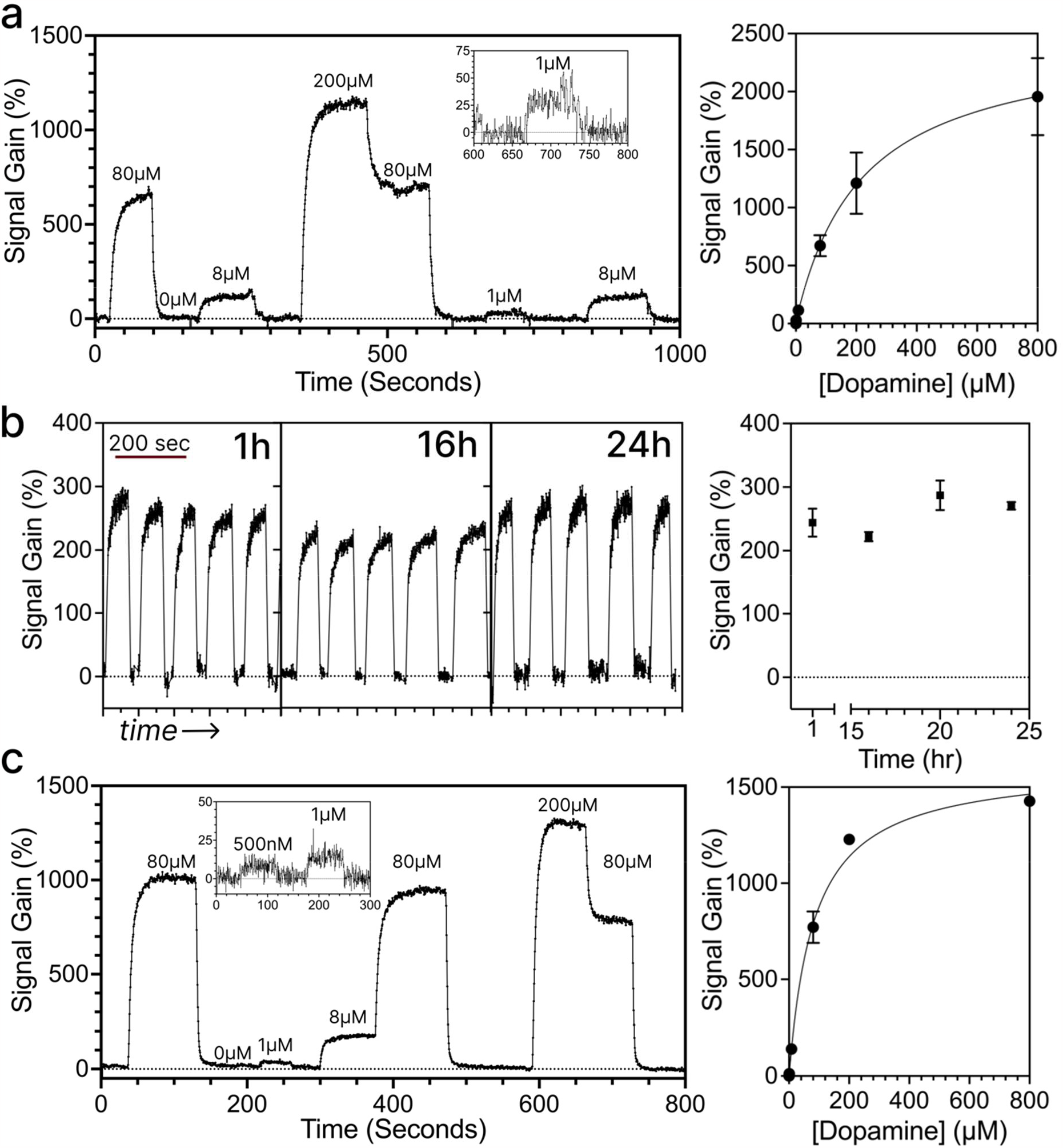
Real-time dopamine detection in buffer and artificial cerebrospinal fluid (aCSF). Continuous, real-time measurement of dopamine in **a)** buffer and **b**) aCSF. Insets show magnification of signal gain produced at 1 µM dopamine in buffer or 500 nM and 1 µM dopamine in aCSF. Righthand panels show standard curves relating signal gain to dopamine concentration. **c)** The sensor maintains consistent response after multiple cycles of dopamine addition and wash after 1, 16, and 24 h. The plot on the right shows the average signal gain distribution of 10 cycles for time-points at 1, 16, 20, and 24 h. Error bars in all panels represent the standard deviation of the average (n = 3). Data fits and normalization are explained in Methods.

Our sensor also maintained high sensitivity over extended periods of time in buffer. We monitored the signal gain in response to multiple sets of 10 cycles of 8 μM dopamine followed by buffer wash (100 s/cycle); we repeated this process at time intervals of 1, 16, 20, and 24 h after the start of the experiment (**Fig. 5b**). Upon the first addition of dopamine at time zero, the sensor produced an average signal gain of 244%. After 16 h, the signal gain from the sensor decreased only slightly to 224%, and even after 24 h, we still observed an average signal gain of 271% over the course of a total of 40 dopamine-buffer cycles (average CV per time point = 0.035) (**Fig. 5b**). Importantly, the sensor consistently returned to baseline with minimal drift (CV = 0.12) in buffer, even after 24 h. We also saw minimal inter-probe variability based on the signal gain produced after applying 8 μM dopamine to three different fiber probes (CV = 0.142) (**Supplementary Figure S13**). In order to test our sensor under more physiologically relevant sensing conditions, we repeated the above experiments in artificial cerebrospinal fluid (aCSF), which features salt and sugar content as well as specific osmolarity and pH that mimic natural CSF in the brain (see Methods). Our sensor achieved essentially comparable performance in aCSF as we observed in buffer (**Fig. 5c, Supplementary Table 6)**, with a dynamic range of 500 nM to 800 μM dopamine and an average CV per concentration of 0.04. We also observed temporal resolution in aCSF (k_on_ = 149.4^−1^s^−1^ and k_off_ = 0.169s^−1^) that was consistent with our results in buffer.

### Continuous detection of cortisol in undiluted human plasma

To demonstrate the generalizability of our platform, we developed a DBS-based sensor for the steroid hormone cortisol. Starting with a previously published cortisol aptamer^27^ (51 nt, K_D_ = 1 µM), we generated multiple DBS switches with variable L_DS_ and L_bubble_ (**Supplementary Table 1**) and tested their affinity in solution using a plate reader-based assay. Decreasing L_bubble_ from 32 to 22 nt while maintaining a constant L_DS_ of 8 nt improved the aptamer affinity by almost 7-fold from 97.46 μM to 14.24 μM. Additionally, as with the dopamine DBS, decreasing L_DS_ while maintaining a constant L_bubble_ resulted in higher sensitivity, with a decrease in K_Deff_ from 97.46 μM for 32-8 to 35.03 μM for 32-7 (**Supplementary Figure S14, Supplementary Table 7**).

We therefore selected 22-8 for coupling to our fiber probe, as this construct achieved a dynamic range (500 nM–175 µM) that fell closest to expected physiological concentrations (50 nM–1 µM).^28,29^ After fabricating the fiber probe as described above, we exposed our sensor to varying cortisol concentrations in buffer (0 nM, 200 nM, 1.4 μM, 10 μM, 140 μM). Our sensor exhibited clear and proportional signal gain across this concentration range, from 5.9% at the LOD of 200 nM to 567% at 140 μM (**Fig. 6a**). We observed minimal variance at each concentration tested (average CV = 0.012). This switch also produced a very rapid, sub-second kinetic response (**Supplementary Figure S15)**. By increasing the incident laser power from 1 μW to 5 μW and thereby improving the signal-to-noise ratio, we achieved a LOD of ∼50 nM (**Supplementary Figure S16)**—roughly three orders of magnitude below the *K*_*D*_ of the DBS (∼14 µM).

**Figure 6:**
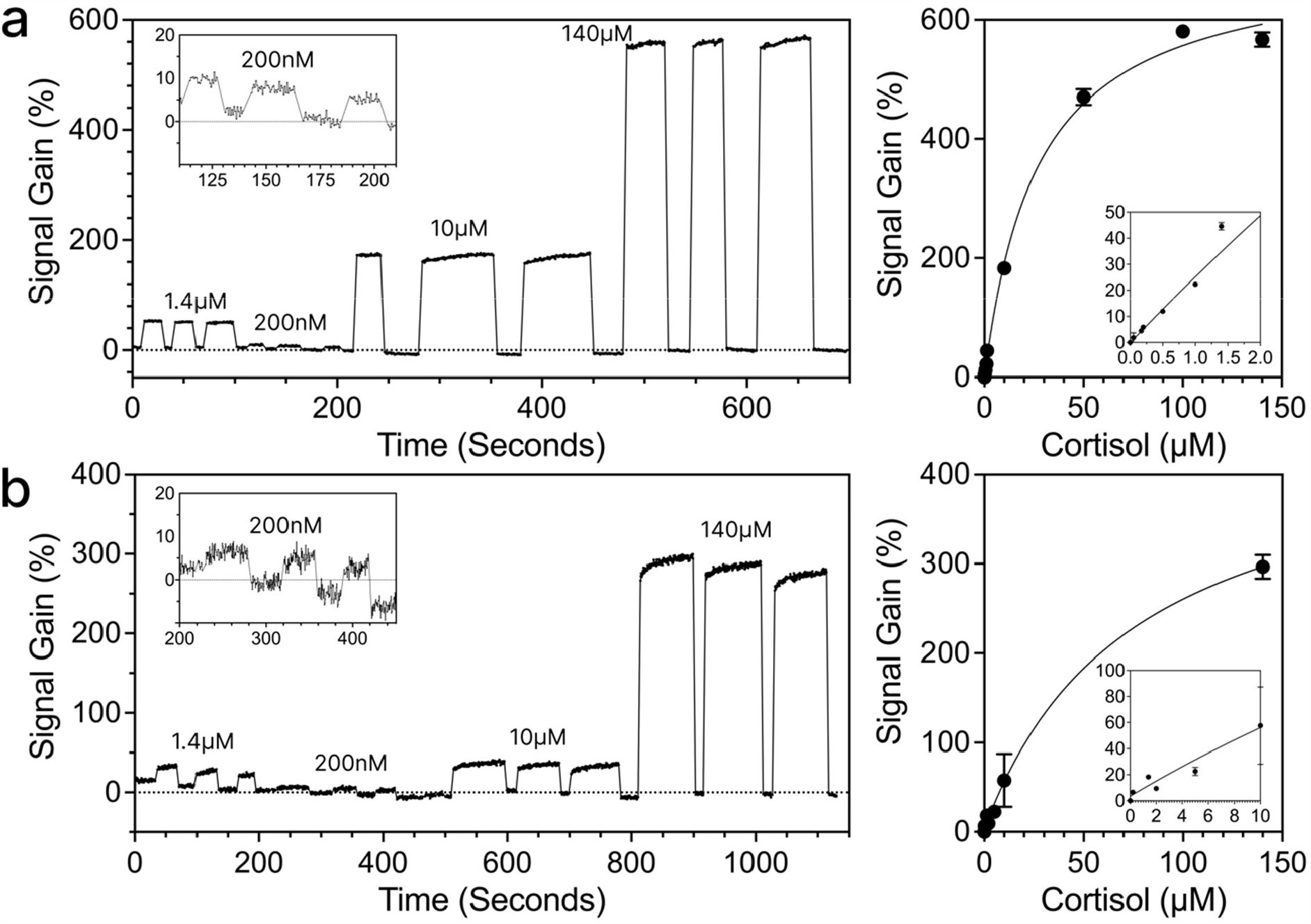
Cortisol detection in buffer and undiluted human plasma. Continuous, real-time measurement of cortisol in **a**, buffer and **b**, undiluted human plasma. Insets show magnification of signal gain at low concentrations, righthand panels show standard curves, with linear ranges presented as insets. Error bars represent the standard deviation of the average (n = 3). Data fits and normalization are explained in Methods.

Finally, we demonstrated that our sensor could achieve robust, continuous cortisol sensing in undiluted human plasma. We sequentially exposed the sensor to varying concentrations of cortisol spiked into plasma (0 μM, 200 nM, 1.4 μM, 10 μM, and 140 μM) and observed close correlation in terms of signal gain (**Fig. 6b**), ranging from 6.5% at 200 nM to 296% at 140 μM, with a similar LOD (200 nM) compared to our buffer measurements. We also observed slightly higher variability in our replicate measurements for each concentration tested on the probe (average CV = 0.05). This increase in CV and decrease in signal gain (**Supplementary Table 6**) may be attributable to a number of reasons, including the fact that free cortisol is known to bind to the corticosteroid-binding proteins globulin and albumin in plasma, potentially resulting in depletion of the spiked-in cortisol in our experiments.^30^ Nevertheless, these results clearly demonstrate that our DBS-coupled probes can achieve rapid, quantitative detection of protein analytes in complex, minimally-processed samples, and highlight the generalizability of this approach.

## CONCLUSION

In this work, we have demonstrated a generalizable platform for continuous, quantitative molecular detection in complex biological samples using structure-switching aptamers with an optical read-out. Our system adapts existing aptamers into a modular DBS construct, which is then coupled to a surface-functionalized fiber-optic probe that detects binding-induced fluorescent signals within an evanescent field, thereby mitigating the confounding effects of surface fouling and autofluorescence from interferents present within the sample. As proof of concept, we generated sensors for two different analytes—dopamine and cortisol—and demonstrated that we could detect nanomolar concentrations of these targets in both buffer and complex specimens like aCSF or undiluted human plasma. Furthermore, our platform could rapidly respond to both increases and decreases in analyte concentration with sub-second resolution, while demonstrating the potential to achieve robust performance over longer periods of use. We saw reasonably consistent signal gain with our dopamine measurements after more than 24 h in buffer (**Fig. 5b**), and even with undiluted plasma, we were able to achieve fairly stable performance over the course of multiple hours (**Supplementary Fig. S17**). Our initial demonstration with cortisol in plasma also hints at the potential clinical utility of this sensor design. The normal range of cortisol concentrations in plasma fluctuates from 136–690 nM in the morning to 55–386 nM in the evening^28,29,31^, but pathological states such as heart disease can give rise to considerably elevated levels of cortisol. Our proof-of-concept sensor showed a similar LOD in plasma relative to buffer (200 nM), which may be sufficient to detect elevated cortisol levels as a biomarker for disease or injury. Although cortisol sensitivity was diminished in plasma, we believe this is principally attributable to the choice of target and the detrimental impact of cortisol-binding proteins present in the matrix. More generally, we observed excellent sensitivity with our sensors, achieving LODs that were typically several orders of magnitude lower than the *K*_*D*_ of the DBS component itself; this suggests that sensors based on even higher-affinity aptamers (*e*.*g*. low nanomolar *K*_*D*_) should be able to monitor even lower-abundance analytes with good precision. And as demonstrated with both analytes, increased laser power can greatly improve the sensor’s sensitivity at the cost of a shorter time window for detection due to heightened photobleaching.

However, there are also clear opportunities to improve this technology in future iterations. For one, the initial aptamer DBS switch design effort currently entails a trial-and-error process in which multiple constructs are synthesized and evaluated. High-throughput screening methods could broaden the scope of this process while also accelerating the identification of switch constructs that offer the best dynamic range and kinetics for a particular target in a given sample matrix. Additionally, although we were able to achieve robust performance in terms of cortisol detection in human plasma for a few hours, the signal gain subsequently degraded over longer periods of time (**Supplementary Figure S17**). It remains unclear at present what contributes to the decline in performance beyond this point. Our system is designed to counter the effects of biofouling, but there are other factors that are likely to contribute to this degeneration over time, including the degradation of the natural DNA-based DBS constructs in the complex plasma environment. In other work, we have shown that the use of nuclease-resistant, chemically-modified nucleobases^32^ can greatly enhance aptamer durability, and this will be worth exploring further in the present sensor design. Sensor performance could also be further enhanced by using alternative fluorophores that minimize the effects of photobleaching and photodegradation, or the implementation of ratiometric-based fluorescence measurements such as FRET, which can contribute to greater sensitivity and lower background. In future work, we intend to optimize this generalizable sensor framework in order to further improve its stability and sensitivity in complex biosamples, with the goal of achieving sensitive longer-term analyte detection in flowing blood and other clinically-relevant sample matrices.

## METHODS

### Reagents

All chemicals were purchased from Thermo Fisher Scientific unless otherwise noted. Oligonucleotides modified with Cy3, Iowa Black, or BHQ-2 were purified by HPLC and purchased from Integrated DNA Technologies. All sequences used in this work are shown in **Supplementary Table 1**. All oligonucleotides were resuspended in nuclease-free water and stored at −20 °C. All experiments were performed in triplicate unless otherwise noted.

### Measurement of effective binding affinity

DBS constructs of varying L_bubble_ and L_DS_ were suspended at a concentration of 250 nM in binding buffer (1X PBS, pH 7.5 and 2 mM MgCl_2_). Switches were prepared by mixing 1 µM Cy3-labeled aptamer strand with 2 µM quencher-coupled strand, heating to 95 °C, and then cooling to 25 °C over the course of 25 minutes to achieve hybridization. To obtain binding curves in solution, 40 µL reactions were prepared in binding buffer with 250 nM DBS and final target concentrations in the range of 1 μM–1 mM for dopamine and 500 nM–175 μM for cortisol. We prepared stock solutions of 20 mM dopamine in 1.5 mL of binding buffer and 30 mM cortisol in 1.5 mL of 100% ethanol. The fluorescence spectra for all samples were measured at 25 °C on a Synergy H1 microplate reader (BioTeK). Emission spectra were monitored in the 550–700 nm range with Cy3 excitation at 530 nm and a gain of 100 in Corning 96-well half area black flat-bottom polystyrene microplates. Representative concentration-dependent emission spectra are shown in **Supplementary Figure S1**.

K_D_^eff^ was extracted from triplicate data by fitting to the single-site specific binding equation in terms of relative fluorescence units (RFU) using GraphPad Prism (**Supplementary Table 2**):

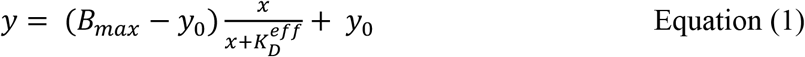

For ease of comparison, normalized data is presented in **Figure 2**. RFU data was first averaged to obtain a mean RFU per concentration point. Averaged data were then normalized as follows, where F_min_ is the minimum mean value and F_max_ is the maximum mean value:

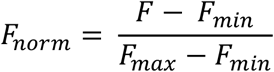

### Measurement of binding kinetics

DBS constructs of varying L_bubble_ and L_DS_ were suspended at a concentration of 333.3 nM in 30 μL of binding buffer. The switches were prepared as described above. Kinetic fluorescence measurements were made using a Synergy H1 microplate reader.

After timed injection of 10 µL of 50 µM dopamine in binding buffer into the 30 μL DBS solution, we measured the kinetic response. Cy3 was excited at 530 nm, and emission was measured at 570 nm using monochromators at the minimum possible regular time interval of 0.465 s. We normalized all kinetic data relative to the target-free control to account for the effect of sample volume change upon injection of dopamine. For plotting, we normalized the curves to a range of 0 to 1 in order to visually emphasize changes in rate constants rather than the background and peak levels that are dictated by the thermodynamics. These data were normalized as described above. Each replicate was normalized independently, and the normalized datasets were then averaged. The association time constant (τ) was obtained by fitting the data to the one-phase association curve using GraphPad Prism (**Supplementary Table 3**).

### Slide preparation for TIRF microscopy

A detailed version of the coverslip cleaning procedure can be found in the Supporting Information of reference [18]. Briefly, coverslips (25 cm x 25 cm x 170 μm) were soaked in piranha solution (25% H_2_O_2_ and 75% concentrated H_2_SO_4_) and sonicated for 90 min, followed by five rinses with Milli-Q water. The coverslips were then soaked with 0.5 M NaOH and sonicated for 30 min, followed by five more rinses with water. The slides were then soaked in HPLC-grade acetone for 5 min while sonicating, and then dried with N_2_.

To prevent nonspecific adsorption of biomolecules onto the glass surface, the coverslips were functionalized prior to use with poly(ethylene glycol) silane (mPEG-sil, MW = 5,000, Laysan Bio). For sample immobilization, a 99:1 ratio w/w of PEG-sil/biotin-PEG-sil was used. The coverslip was incubated with a 25% (w/w) mixture of PEG-sil (or PEG-sil/biotin-PEG-sil) in DMSO (anhydrous) at 90 °C for 15 min. The excess PEG was rinsed off the coverslip with water, and the coverslips were dried under an N_2_ stream. Imaging chambers (∼8 μL, 10 mm x 20 mm x 0.25 mm) were constructed by pressing a polycarbonate film with an adhesive gasket (Grace Bio labs) onto the PEG-coated coverslip. Before image acquisition, the surface was incubated with 12 μL of 0.2 mg/mL (∼200 nM) streptavidin solution for 10 min. The unbound streptavidin was washed away twice with 50 μL of T50 imaging buffer (50 mM NaCl, 20 mM Tris buffer pH 8.0). For surface attachment, the displacement strand was conjugated at its 3’ end to a biotin moiety and at its 5’ end to an Iowa Black quencher, whereas the aptamer complement was internally modified with a Cy3 reporter. These DBS switches were assembled as described above, and then diluted in binding buffer to the desired final concentration. We then incubated the passivated, streptavidin-coated coverslips with 250 nM DBS in 1x binding buffer for 1 h at room temperature, followed by washing with more binding buffer.

### TIRF microscopy and image analysis

Fluorescence imaging was carried out using an inverted Nikon Eclipse Ti2 microscope equipped with the perfect focus system and implementing an objective-type TIRF configuration with a motorized Nikon TIRF illuminator (LAPP) and an oil-immersion objective (CFI Apo TIRF 60× oil immersion objective lens, numerical aperture 1.49). The effective pixel size was 180 nm. With these settings, a single 532-nm laser was used for excitation (LUN-F XL 532/561/640 Laser Combiner; 20 mW, measured out of the objective). For Cy3 imaging, the laser beam was passed through a multiband cleanup filter (ZET532/640x, Chroma Technology) and coupled into the microscope objective using a multiband beam splitter (ZT532/640rpc-uf2, Chroma Technology). Fluorescence light was spectrally filtered with a ZET532/640m-trf (Chroma Technology) filter. Fluorescence was further filtered with an ET585/65m (Chroma Technology) emission filter mounted on a Ti2-P-FWB-E motorized barrier filter wheel. All movies were recorded onto a 704 × 704 pixel region of a back-illuminated Scientific CMOS camera (Prime 95B, 1.44 MP, Teledyne Photometrics). The camera and microscope were controlled using the NIS-Elements Advanced Research software package. Fluorescence intensity-time trajectories of individual molecules were extracted from the videos using a custom algorithm written in MATLAB.

### Fiber probe selection and construction

Single- and multi-mode fibers are commercially available and have low attenuation (*i*.*e*., ∼3-5 dB/km at wavelengths of 650-850 nm, which is the bandwidth at which Cy3 emits). Background fluorescence was also a consideration; this is likely to originate from impurities in the fiber (color centers) and can dominate the spectrum in a poorly chosen fiber. To minimize this background emission, we screened different types of fibers, including a) single- and multi-mode fibers, b) multi-mode fibers with different core diameters (100, 62.5, or 50 μm), c) multi-mode fibers with step index and graded index, and d) multi-mode fibers with high and low OH content. **Supplementary Figure S7** shows representative transmission spectra for different multimode fibers excited at 405/530/630 nm using appropriate laser lines with band-pass and long-pass filters in the excitation and detection paths. To minimize ambient light coupling to the fiber, which introduces background signal offset, and to prevent damage to the fiber, we use reinforced tubing (Thorlabs FT038-BK) for multimode fibers. The background was lowest when using a graded index multi-mode fiber (Thorlabs GIF625) with a core diameter of 62.5 μm, cladding diameter of 125 μm, and an operating wavelength longer than 550 nm. Since the residual background counts from the fiber are constant, they can be easily subtracted from the measured signal to obtain the Cy3 signal.

Optical fibers confine light in a high refractive index core via total internal reflection, and a radially decaying evanescent field exists in the surrounding medium of lower refractive index. Tapering a fiber to the proper range of core diameter allows for the enhancement of the evanescent field in this region, thereby increasing the light coupled to the aptamer switches grafted on the surface of the taper. This in turn increases the sensing area compared to a fiber sensor with a cleaved end, where coupling can only occur at the fiber core. Evanescent coupling along the fiber taper has the added benefit of rejecting background signal from autofluorescence in biological settings (**Supplementary Figure S6**). In order to finely control fiber geometry and reproducibly fabricate high-performing fibers, we adapted a previously established heat-and-pull-based tapering protocol26. Several inches of plastic coating were stripped from a central section of the fiber before it was tapered using a Vytran GPX3000, using a custom recipe that produced approximately 10-µm fiber waists, with intended up- and down-taper lengths of 10 mm. The taper parameters (i.e. up- and down-taper length and waist diameter) were chosen to produce the best compromise between requirements, namely (1) maximizing the evanescent excitation of the fluorophores, (2) maximizing the selective collection of fluorophore emissions by the fiber core, and (3) reducing non-adiabatic scattering losses. After tapering, the two fiber ends were gently pulled mechanically using the micro-stage in order to cleave the tapered fiber at its waist and obtain two different fiber probes. Fiber tapering and cleavage consistently resulted in two visually distinct profiles (one on either side of the filament) with different taper lengths, referred to as “short” and “long” tapers. Though both tapers were usable, under the same conditions, long tapers generally produced about twice the signal as short tapers and were therefore used throughout the main text (**Supplementary Figure S8**). Our tapering process yielded reproducible results, and while tapers can break, we found them to be mechanically strong enough to not be prohibitively sensitive during our measurements.

### Preparation of the fiber tips

We used a custom-built micro-stage setup to fine tune the functionalized region of the fiber and consistently achieve a reproducible probe coating. The surfaces of the fiber tips were first cleaned and hydroxyl-activated by immersing them in piranha solution for 2 h, followed by a 10 min incubation/wash with ultrapure water, followed by acetone for 5 min. For the amination of the optical fiber, the probes were then kept for 5 min in 1% (v/v) Vectabond/acetone, washed with ultrapure water, and dried for 5 min at room temperature. In order to prevent non-specific adsorption of biomolecules onto the glass surface, these surface-attached amine groups were incubated with a mixture of poly(ethylene glycol) succinimidyl valerate (mPEG-SVA, MW = 5,000) and biotin-PEG-SVA at a ratio of 90:10 (w/w) in 110 mM sodium bicarbonate for 3 h and then rinsed with water and left in a dried state until used.

A polycarbonate chamber (∼40 μL, Grace Bio labs HybriWell Sealing systems) was then constructed on the probe tip by pressing a polycarbonate film with an adhesive gasket onto the coated fiber in order to surround the probe (**Supplementary Figure S5**). Two silicone connectors were glued onto the predrilled holes of the film and served as inlet and outlet ports to manually dispense/wash different solutions into the chamber. Before data acquisition, the chamber was filled with 40 μL of 0.2 mg/mL (∼200 nM) neutravidin solution for 10 min. The unbound neutravidin was washed away twice with 200 μL of 1x buffer (1X PBS, pH 7.5 and 2 mM MgCl_2_). Biotinylated Cy3- and quencher-modified DBS constructs were then prepared as described above and immobilized onto the passivated, neutravidin-coated fiber surface by incubating the chamber for 1 h with 40 μL of 250 nM DBS solution in 1x binding buffer followed by washing with more binding buffer. All experiments were conducted at room temperature. In our control experiments, we incubated the PEGylated fiber sensor with 40 μL of 250 nM Cy3-labelled DNA solution in 1x binding buffer in the absence of neutravidin. Measurements were taken with the Cy3-labelled DNA still in solution (solution signal) and after washing the fiber with 1x binding buffer (non-specific binding signal). These measurements were compared to the specific signal after incubating the PEGylated fiber sensor with neutravidin, washing, incubating with 250 nM Cy3-labelled DNA, and washing (as described above). We observed minimal signal in this control experiment, confirming that 1) there is minimal non-specific binding of aptamer switches to the sensor surface, and 2) the fiber operates as a near-field sensor and only detects signals close to the surface, with minimal interference from fluorophores in solution (**Supplementary Figure S6**).

### Preparation of target solutions for real-time measurements

For the dopamine experiments, we prepared a stock of 20 mM dopamine in 1.5 mL of binding buffer or aCSF (124 mM NaCl, 2.5 mM KCl, 10 mM glucose, 26 mM NaHCO_3_, 1 mM NaH_2_PO_4_, 2.5 mM CaCl_2_, and 1.3 mM MgSO_4_)^33^. We then performed serial dilutions to prepare solutions of 0, 200nM, 500nM 1 μM, 5 μM, 8 μM, 15 μM, 20 μM, 80 μM, 200 μM, or 800 μM dopamine in buffer or aCSF. Because of the issue of polydopamine formation over time in high concentration stocks, we renewed the dopamine stock every 45 min to 1 h to ensure minimal precipitation on the probe surface or fouling of aptamer structures.

For cortisol experiments, since cortisol is poorly soluble in aqueous conditions, we first prepared a stock solution of 30 mM cortisol in 1.5 mL of 100% ethanol. We then did a serial dilution in binding buffer to prepare solutions of 50 nM, 200 nM, 500nM, 1 μM, 1.4 μM, 10 μM, 50 μM, 100 μM, 140 μM of cortisol. We subsequently injected different solutions into the sample chamber, exchanging solutions at a 1-min interval while continuously recording. For experiments with human plasma, we used the cortisol stock in 100% ethanol to prepare serial dilutions in human plasma at 99.5% final plasma concentration (50 nM, 200 nM, 500nM, 1 μM, 1.4 μM, 50 μM, 140 μM). It is important to note that the dilutions for each sample were freshly prepared right before measurements because cortisol-binding proteins in the plasma samples contributed to discrepancies in measurements when we used older, pre-prepared samples. Source data for all concentrations tested are provided as a Source Data file.

### Optoelectronic system set-up for real-time measurements

Fluorescence detection was carried out using a single-photon counting module (SPCM), which was interfaced to a custom-designed acquisition board based on an Arduino to process and analyze detected photons, record detection events, and monitor fluorescence intensity over time. Data acquisition was performed with a LabVIEW GUI with an integration time of 500 ms. For excitation, we used a fiber-coupled 532 nm laser (Coherent, 20 mW, measured out of the fiber). For Cy3 measurements, the laser beam is passed through a band-pass cleanup filter (532/10, Chroma Technology) and variable neutral density filter (Thorlabs NDC-50C-4M) and coupled into the sample fiber using a fiber coupler (62.5/125 µm multimode fiber coupler 90/10, FC/PC, Thorlabs) and fiber switch (Newport MPSN-62-12-FCPC) using the appropriate optics and mounts (In-Line Fiber Optic Filter Mount, FC/PC, Thorlabs). The fiber switch modulated the on/off state of the laser beam, allowing the precise control of the duration and timing of the laser pulse for continuous and intermittent measurements. Fluorescence emission light from the fiber sample is collected back through a fiber coupler (62.5/125 µm multimode fiber coupler 50/50, FC/PC, Thorlabs) and spectrally filtered with band-pass filters (595/50, Chroma Technology; 676/29, Semrock). Aligning the fluorescence emission signal with the SPCM using lenses (ø8mm Achromatic Doublet, Threaded mount, Thorlabs) and mirrors is critical to ensure that the incident light is focused, well-centered, and collimated on the active area of the detector, allowing the highest detection efficiency while rejecting other background light.

Data collected from the SPCM was analyzed in MATLAB. Background signal was measured for each fiber probe (laser power at 500 nW, no DBS on the probe surface) and averaged to calculate a background value per fiber. Data was first segmented into sections of different concentrations either by finding the edges of regions of 0 s (for intermittent data with SPCM turned OFF between measurements) or manually (for continuous data). The signal per concentration section was calculated in order to extract a binding curve. If the data increased or decreased by more than 5% over the section, the data was fitted to an exponential gain or decay, and the plateau value was reported. If the decreased by more than 15% over the section, the data was fitted to an exponential decay, and the plateau value was reported. The exponential rate was also returned and used to characterize kinetics of the DBS constructs on fiber. Otherwise, the average value over the section was calculated. The background value of the fiber was subtracted from all calculated signals. Baseline values were defined as the average background-corrected signal over times when no target was present. Signal gain values reported in binding curves were calculated by normalizing the background-corrected signals to the average of the nearest baseline sections prior to and after the concentration was measured. Real-time data presented was normalized by the average baseline value over the entire data file. Signal gain values reported in **Figure 5b** were normalized by the average baseline within the 5 (or 10) cycle measurements. In these experiments, LODs were defined as the lowest experimentally measured concentration for which the average signal gain from triplicate experiments was significantly different from the average background signal gain values in the adjacent baseline segments that preceded sample addition (P < 0.05 on an unpaired t-test).

## Acknowledgements

We would like to acknowledge the contribution of Michael Silvernagel, Xiaoyuan (Sandra) Hu, Peter Mage, Dong-Wook Park, Anping Li, Alex Codik, Kang Yong Loh, Brandon Wilson, Alexandra Rangel, Sharon Newman, Dries Vercruysse and all current members of the Soh lab for their important help throughout the project. We would also like to acknowledge the contribution of the members of the Malenka lab, including Wade K. Morishita, Daniel Cardozo Pinto, Mathew B. Pomrenze, Paul Holbert and Prof. Robert Malenka, who provided valuable insights, productive discussions, and experimental help throughout the course of the work. We would also like to thank SNF and SNSF for providing access to cleanroom and SEM facilities. H.T.S would like to acknowledge support from the Helmsley Trust, Wellcome LEAP SAVE program, and the National Institutes of Health (NIH, OT2OD025342). A.A.H. acknowledges support from the Sanjiv Sam Gambhir—Philips Fellowship Program in Precision Health and the NSERC Postdoctoral Fellowships (PDF, Canada). C.D. acknowledges support from the Andreas Bechtolsheim Stanford Graduate Fellowship and the Microsoft Research Ph.D. Fellowship. A.P.C. acknowledges support from the NSF Graduate Research Fellowship Program and the Stanford Graduate Fellowship. S.Y. acknowledges support from the Stanford Graduate Fellowship program. I.A.P.T. acknowledges the support of the Medtronic Foundation Stanford Graduate Fellowship and the Natural Sciences and Engineering Research Council of Canada (NSERC).

## Contribution

H.T.S., J.V., A.A.H., C.D. and A.P.C. conceived the initial concept. A.A.H., C.D. and A.P.C. designed the experiments. A.A.H. designed and optimized the DBS construct. A.A.H., A.P.C., K.F., D.W. and T.F. conducted troubleshooting of aptamer switch designs. B.E.Y. synthesized quencher-labelled DNA. A.A.H. and A.P.C. executed the solution-based experiments. C.D. and A.P.C. designed/fabricated/aligned the optoelectronic system. N.M. designed and fabricated the Arduino board. A.A.H., A.P.C., C.D., and S.Y. executed the fiber-based experiments. C.D. and A.P.C. developed the software and analyzed the fiber-based data. Y.G. designed and executed the single-molecule experiments and analyzed the corresponding data. A.P.C., K.Y., I.A.P.T., B.H.A. and M.D. designed the fiber-tapering protocol. A.P.C. optimized and executed the fiber-tapering protocol. A.A.H., A.P.C., M.E., and H.T.S. wrote the manuscript. All authors edited, discussed, and approved the whole paper.

