## Supplementary Information for "Continuous optical detection of small-molecule analytes in complex biomatrices"

Amani A. Hariri *et al.*

### OUTLINE

**Supplementary Figure S1** | Raw fluorescence spectra from multiple DBS constructs for dopamine
**Supplementary Figure S2** | Binding curves for all DBS constructs for dopamine
**Supplementary Figure S3** | Summary of kinetic results for dopamine
**Supplementary Figure S4** | Single molecule images and traces from the 22-7 DBS construct
**Supplementary Figure S5** | Fiber in flow-through chamber for real-time measurements
**Supplementary Figure S6** | Near-field detection and non-specific binding characterization
**Supplementary Figure S7** | Data treatment for fibers to eliminate background
**Supplementary Figure S8** | Fiber tapering and signal reproducibility
**Supplementary Figure S9** | Photobleaching rates at different laser power
**Supplementary Figure S10** | Data processing and data smoothing
**Supplementary Figure S11** | Additional data for dopamine real-time detection in buffer at higher laser power to improve SNR.
**Supplementary Figure S12** | Kinetics of the dopamine probe sensor in buffer and aCSF
**Supplementary Figure S13** | Fiber-to-fiber signal gain variability for dopamine
**Supplementary Figure S14** | Raw selected analysis of selected cortisol DBS constructs
**Supplementary Figure S15** | Kinetics fits from cortisol real-time measurements.
**Supplementary Figure S16** | Raw data of real-time cortisol detection at ~1–5 μW laser power
**Supplementary Figure S17** | Signal enhancement after undiluted plasma exposure for up to 26 hours
**Supplementary Table 1** | DNA sequences used in this work
**Supplementary Table 2–3** | Effect of $L_{Bubble}$ and $L_{DS}$ lengths on binding affinity and kinetics
**Supplementary Table 4** | Signal gain from dopamine fiber measurements in buffer
**Supplementary Table 5** | Kinetics fits for dopamine fiber measurements in buffer
**Supplementary Table 6** | Signal gain of dopamine and cortisol on fiber measurements in aCSF or plasma
**Supplementary Table 7** | Effect of $L_{Bubble}$ and $L_{DS}$ on effective cortisol binding affinity
**Supplementary Note 1** | Mathematical model and derivation

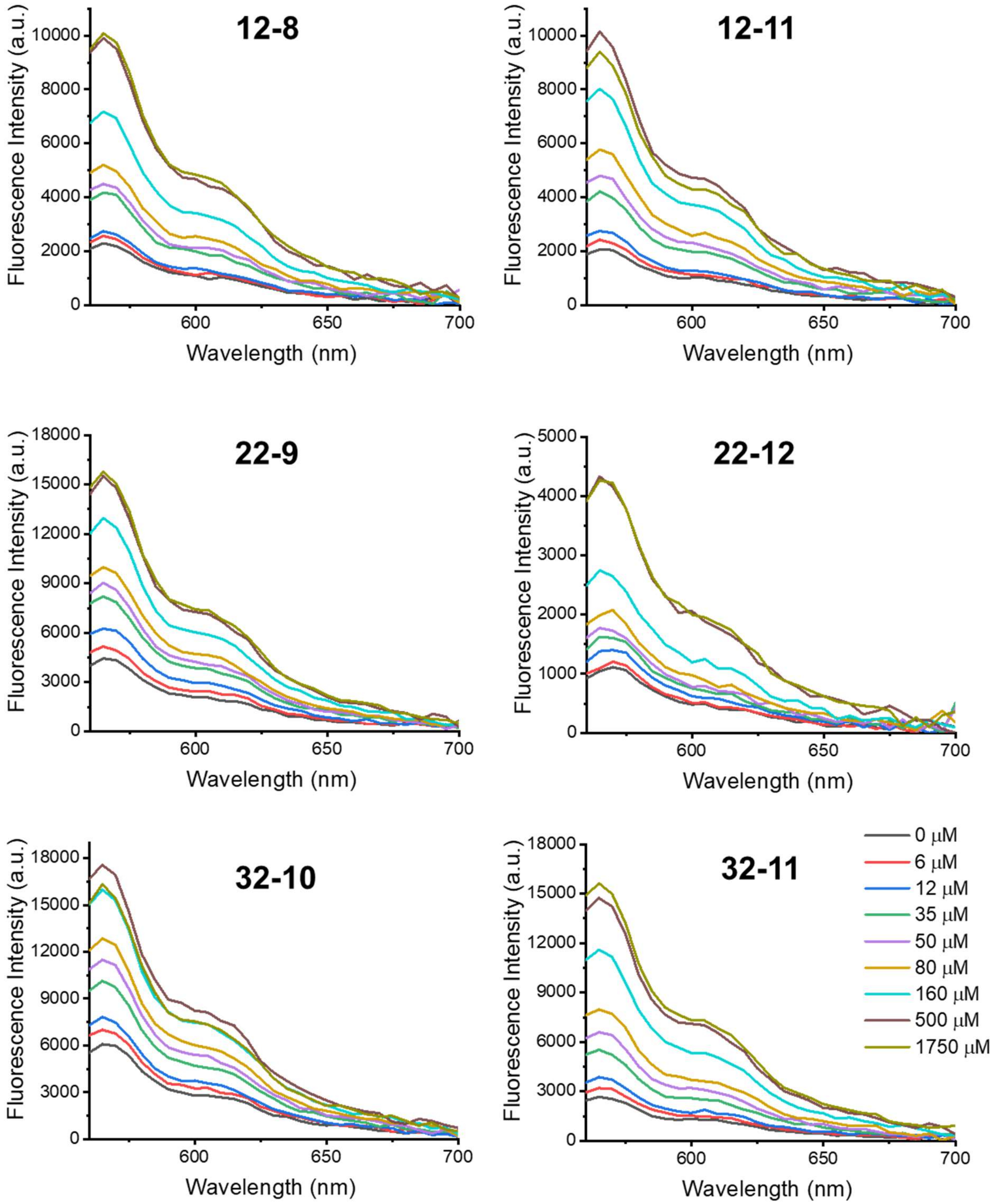

**Supplementary Figure S1: Raw signal analysis of selected dopamine DBS constructs.** Representative concentration-dependent emission spectra of various dopamine DBS constructs. The fluorescence at peak emission (565 nm) was used as raw signal for both thermodynamic and kinetic studies. Colors indicate concentrations of dopamine. All plots show averages over three replicates. Source data for all constructs are provided as a Source Data file.

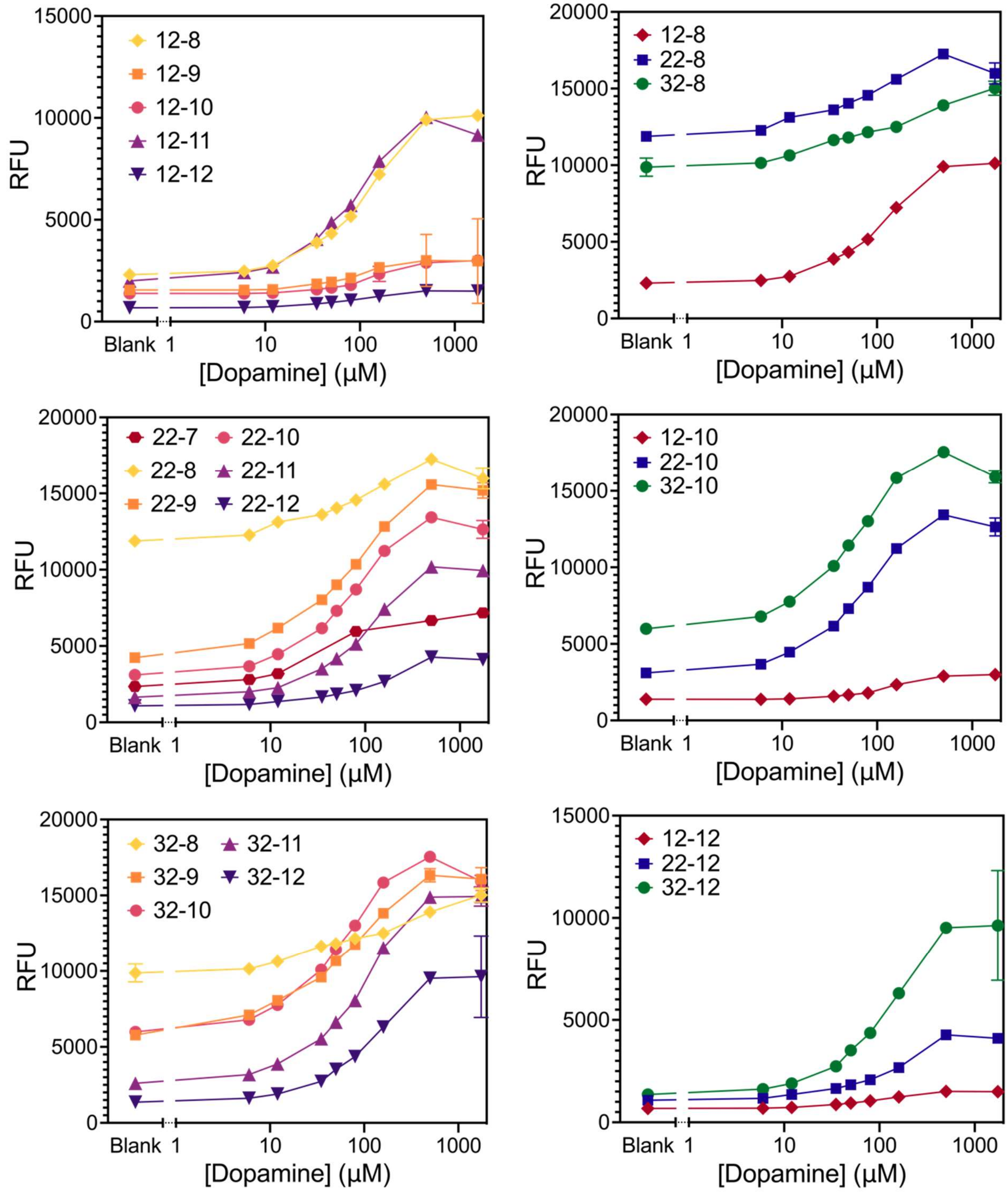

**Supplementary Figure S2: Binding curves for all dopamine DBS constructs.** Plots show average raw fluorescence as a function of dopamine concentration from three replicates. Error bars represent the standard deviation. Source data for all constructs are provided as a Source Data file.

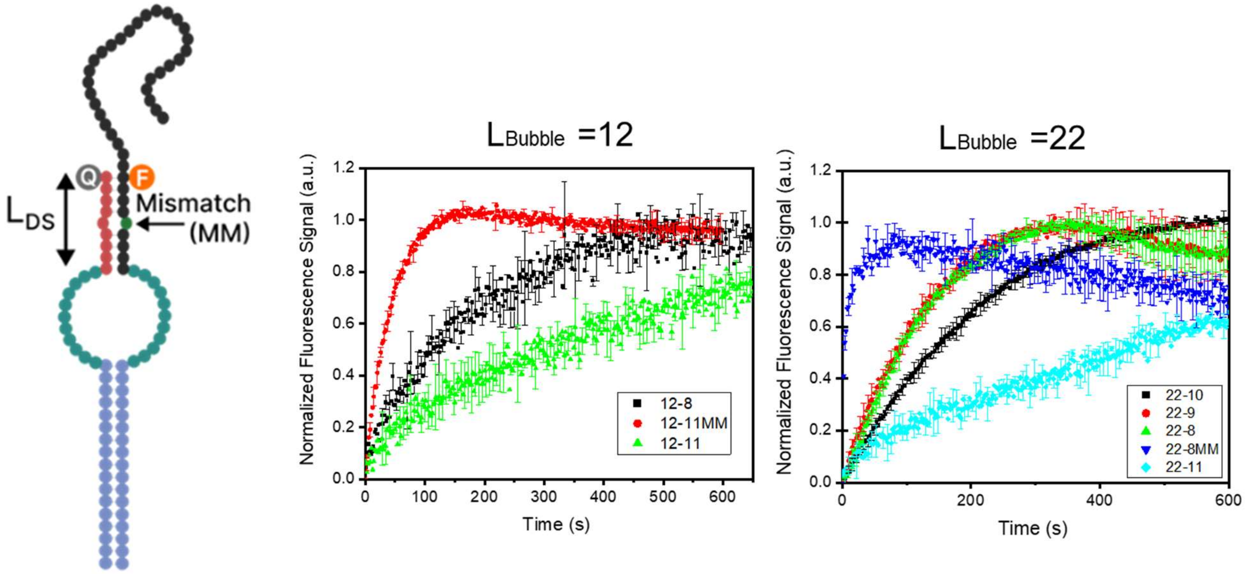

**Supplementary Figure S3: Temporal response modulation via DBS design.** Plots show normalized fluorescence signal upon injection of 50 μM dopamine as a function of time for different DBS constructs. While $L_{Bubble}$ has minimal effect on kinetics, we observed that increasing $L_{DS}$ from 8 to 11 while holding $L_{Bubble}$ constant at 12 nt and 22 results in slower kinetics in all three respective cases. Introducing mismatches greatly increases the signaling kinetics of the constructs, e.g. 12-11 and 12-11MM, where a 11-mer DS with a single mismatch (red) exhibits much faster kinetics faster than the fully complementary 11-mer. All plots are averaged over n = 3 replicates. Error bars represent the standard deviation.

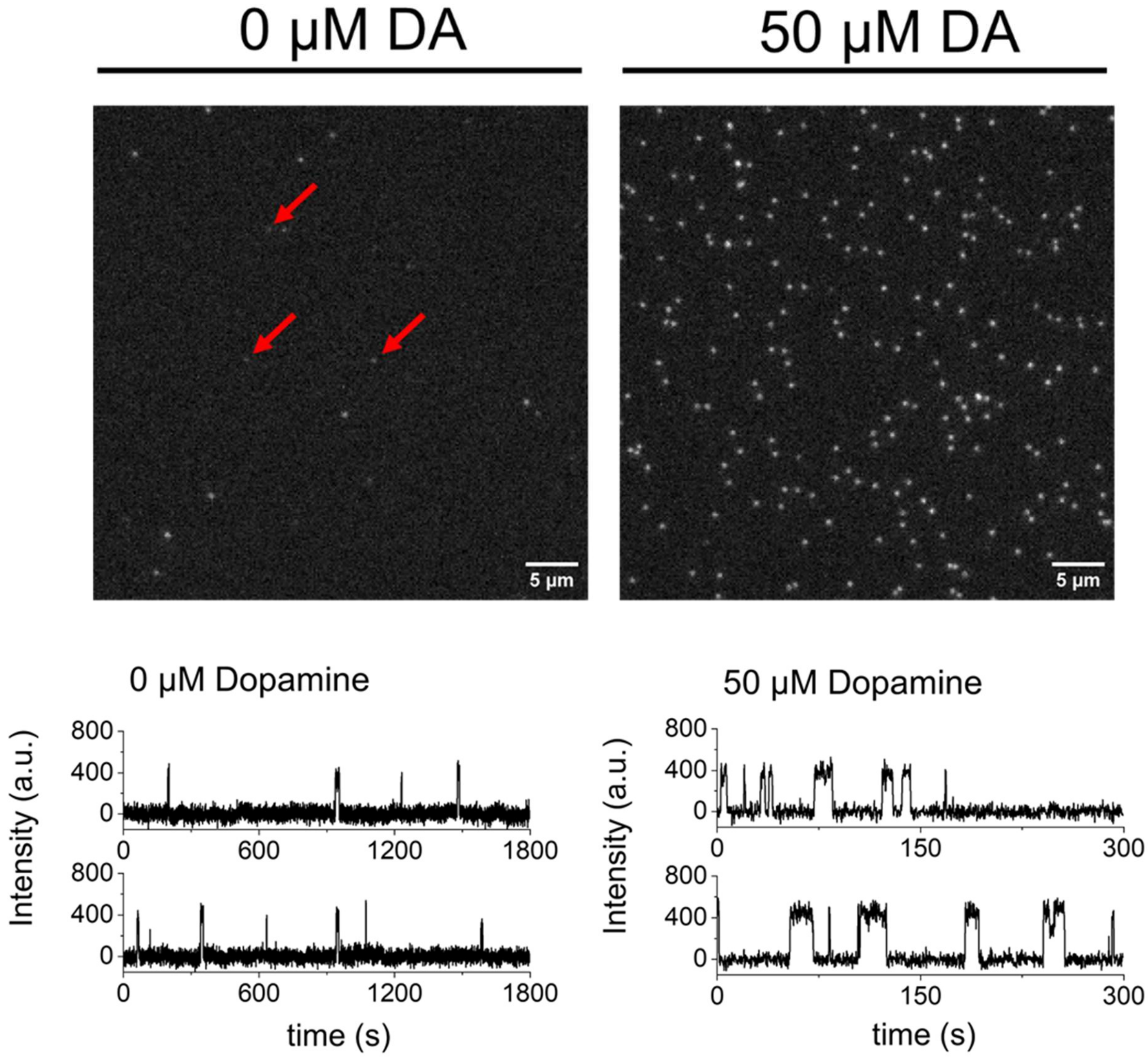

**Supplementary Figure S4: Single-molecule images and traces from the 22-7 DBS construct.** Top panels show fluorescence images of different regions of the coverslip before and after the addition of 50 μM dopamine. Red arrows indicate fiducial markers used for drift correction. Bottom panels show representative intensity-time trajectories. Source data for all traces are provided as a Source Data file.

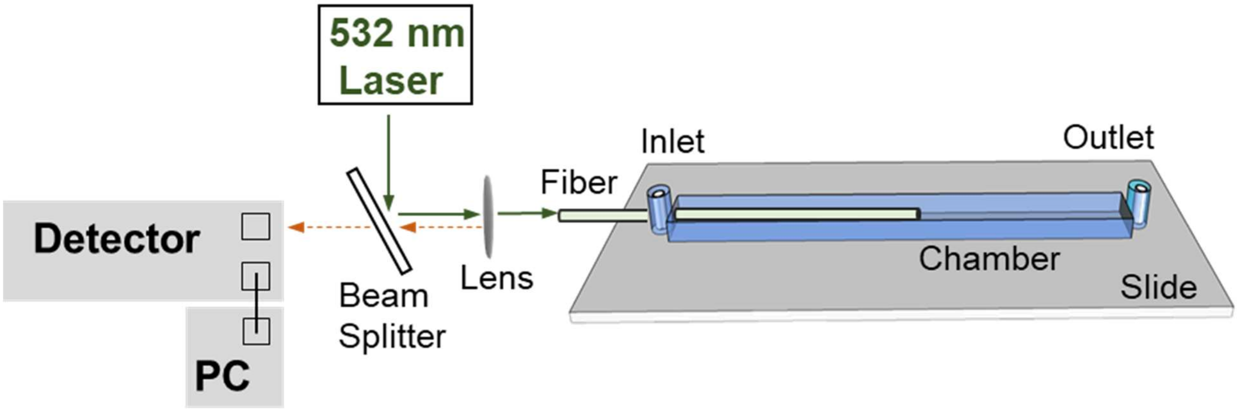

**Supplementary Figure S5: Flow-through chamber setup for real-time measurements.** A polycarbonate chamber was constructed around the probe tip by pressing a polycarbonate film with an adhesive gasket onto the coated fiber. Two silicone connectors were glued onto the predrilled holes of the film and served as inlet and outlet ports to manually dispense/wash different solutions into the chamber. Diagram is not drawn to scale.

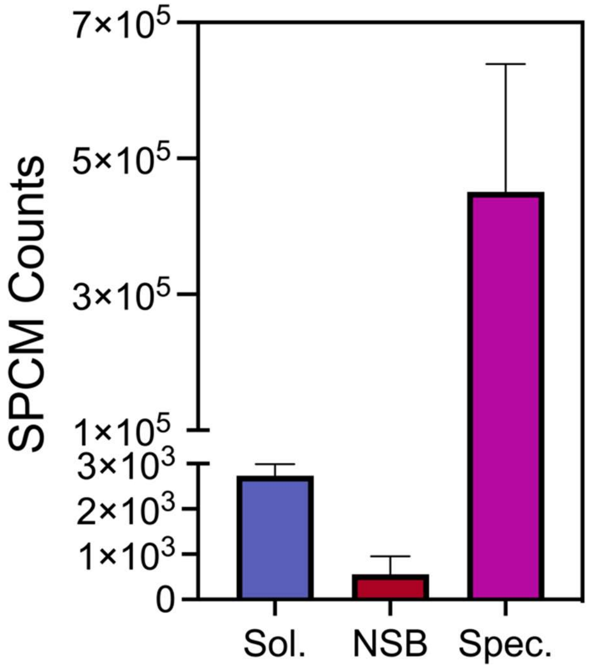

**Supplementary Figure S6: Near-field detection and non-specific binding characterization.** We collected control measurements from PEGylated fiber sensors incubated with 40 μL of 250 nM Cy3-labelled DNA in 1x binding buffer in the absence of neutravidin (Sol.; blue bar) to characterize near-field efficiency in rejecting solution-phase fluorescence signal, followed by additional measurements after washing the fiber to measure the extent of non-specific binding signal (NSB; red bar). These measurements were compared to the specific signal (Spec.; purple bar) derived after following the full functionalization protocol as described in Methods in the presence of neutravidin. Bars depict aggregate data from two short tapers and two long tapers. Error bars represent the standard deviation (n=4).

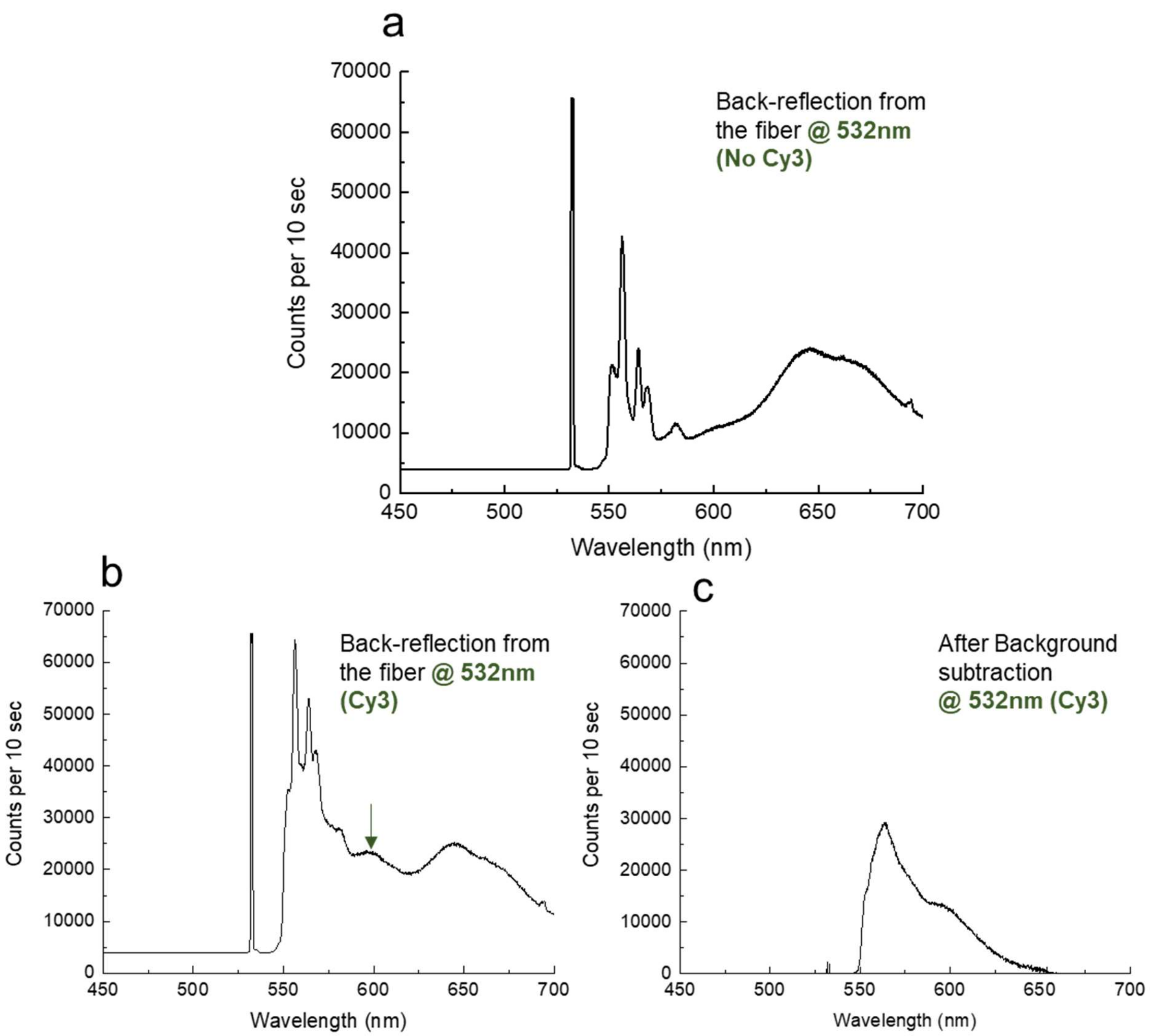

**Supplementary Figure S7: Data treatment for fibers to eliminate background.** Intensity traces from fiber GIF625 (Thorlabs) showing **A)** the background in the absence of Cy3 sample, **B)** signal in the presence of Cy3 sample, and **C)** Cy3 sample signal after background subtraction. GIF625 had the lowest background signal of all fibers tested.

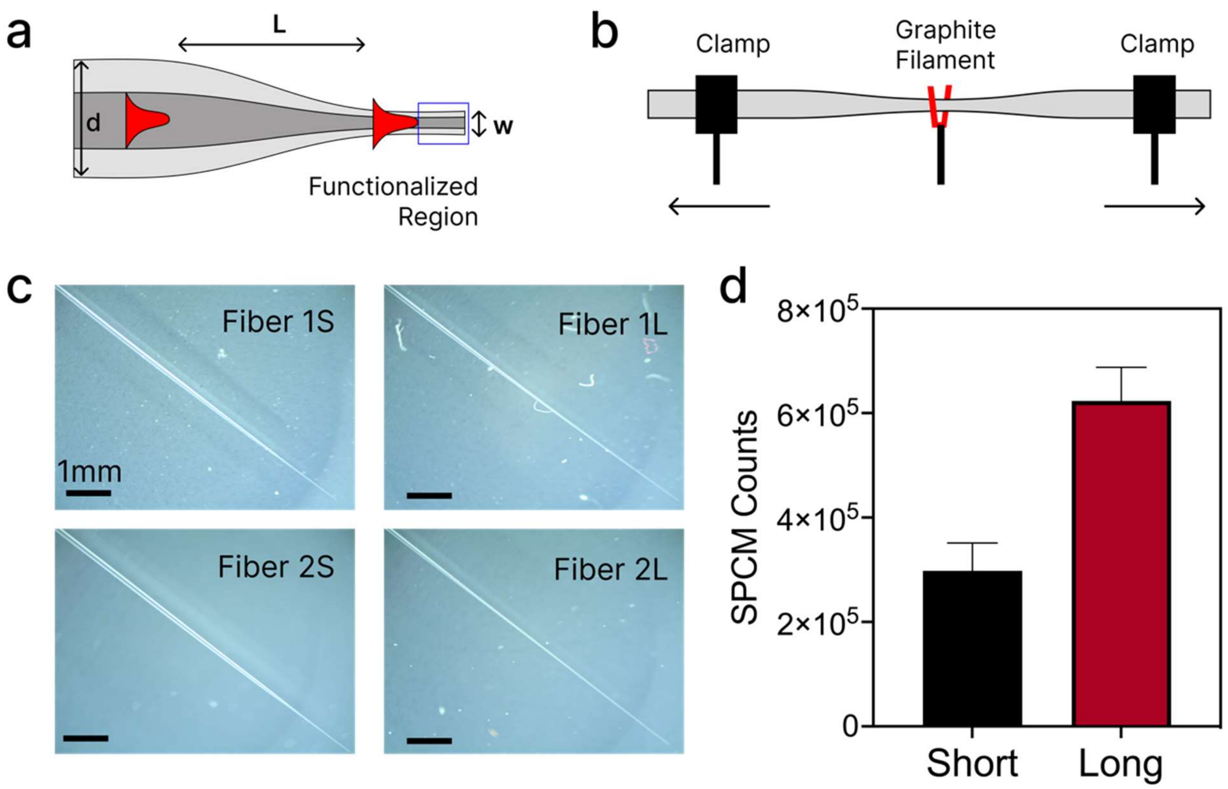

**Supplementary Figure S8: Fiber tapering and signal reproducibility. a)** Scheme of the tapered fiber and optical modes within the unfunctionalized versus functionalized regions. Input recipe parameters for fiber pulling were initial fiber diameter (d) = 125 µm, taper length (L) = 10 mm, and final waist diameter (w) = 10 µm. **b)** Heat-and-pull-based tapering set-up using a Vytran GPX3000. Several inches of plastic coating of the fiber are stripped from a central section of the fiber before it is tapered. Fiber ends are gently pulled mechanically using the micro-stage in order to separate the tapered fiber at the waist into two different fiber probes. **c**) Microscope images showing the two resulting tapered fiber profiles from the pulling technique (L = long, S = short). Scale bar = 1 mm. **d)** Histogram of average SPCM counts from short (N = 2) and long (N = 3) fiber tapers functionalized with 250 nM Cy3-labelled DNA measured at 500 nW incident laser power. Long tapers consistently produced approximately double the signal of short tapers, and the CV of signal produced by long taper was 0.10.

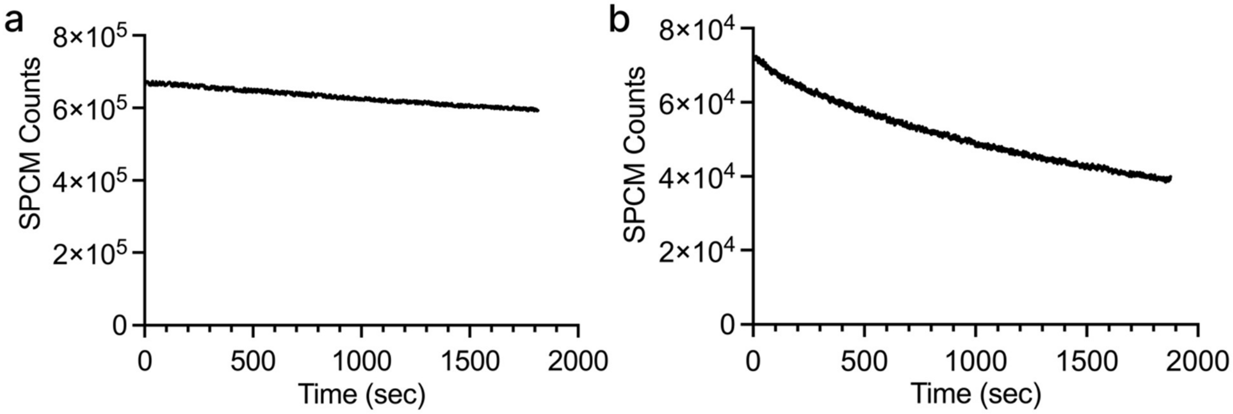

| Power ($\mu$W) | Average τ (sec) | CV |
|---|---|---|
| 0.5 | 5,103 | 0.049 |
| 5 | 1,128 | 0.181 |

**Supplementary Figure S9: Photobleaching rates at different levels of laser power.** Photobleaching of Cy3-tagged DNA sample at **a)** 500 nW or **b)** 5 $\mu$W. SPCM counts are lower in the 5 $\mu$W experiment due to neutral density filters positioned in front of the SPCM to prevent detector damage due to the increased photon number. The table summarizes the average single exponential decay time constant (τ) (n = 3).

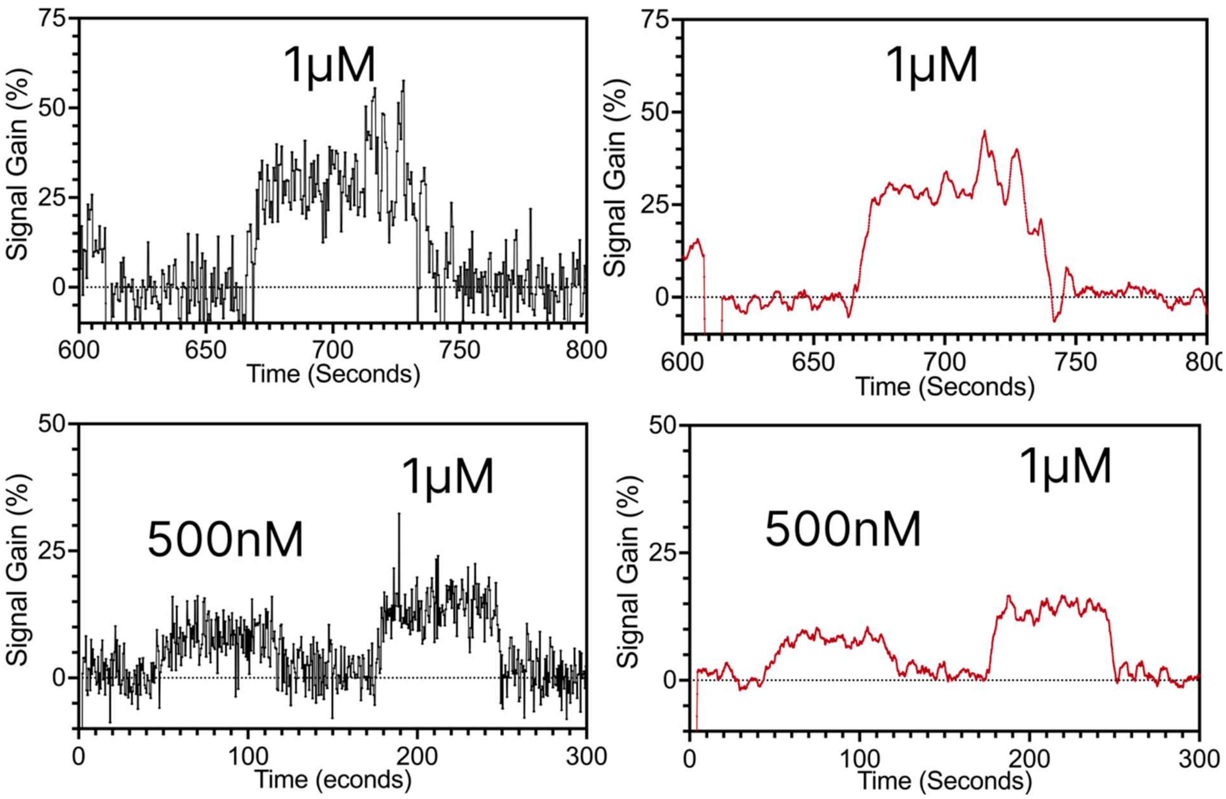

**Supplementary Figure S10: Data processing and data smoothing.** Comparative plot showing non-smoothed (left) and data smoothed with a N = 50 moving average filter in MATLAB (right). Data shown are the insets from **Fig 5a, b**.

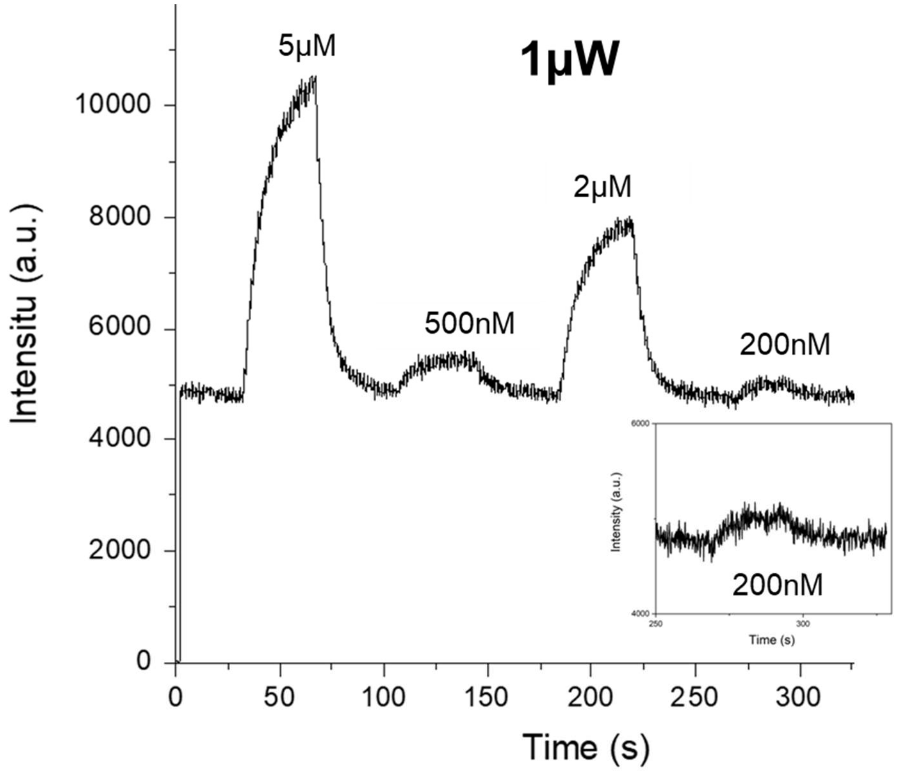

**Supplementary Figure S11: Additional data for dopamine real-time detection in buffer at higher laser power to improve SNR.** Raw data showing real-time dopamine measurements in buffer at 1 µW laser power.

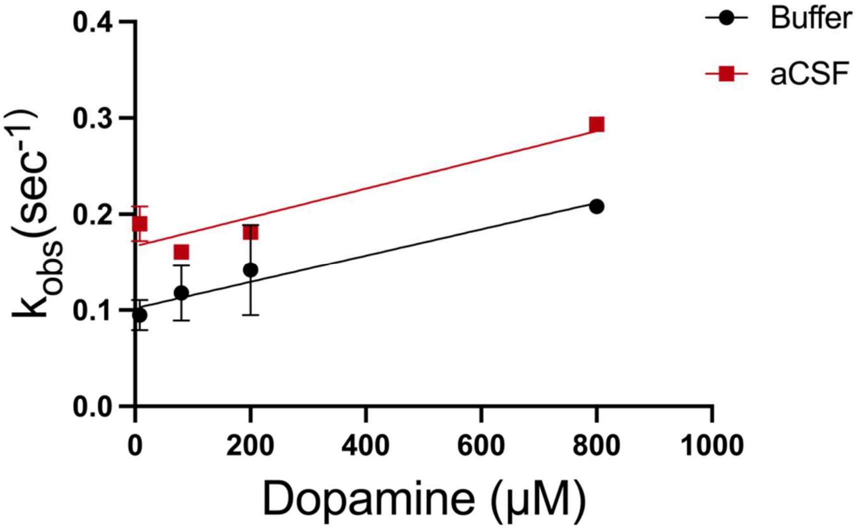

**Supplementary Figure S12: Kinetics of the dopamine probe sensor in buffer (black) and aCSF (red).** $k_{obs}$ values were extracted from real-time data in MATLAB. $k_{on}$ and $k_{off}$ were extracted using linear regression in GraphPad Prism with the following equation ($k_{obs} = k_{on}[Dopamine] + k_{off}$).

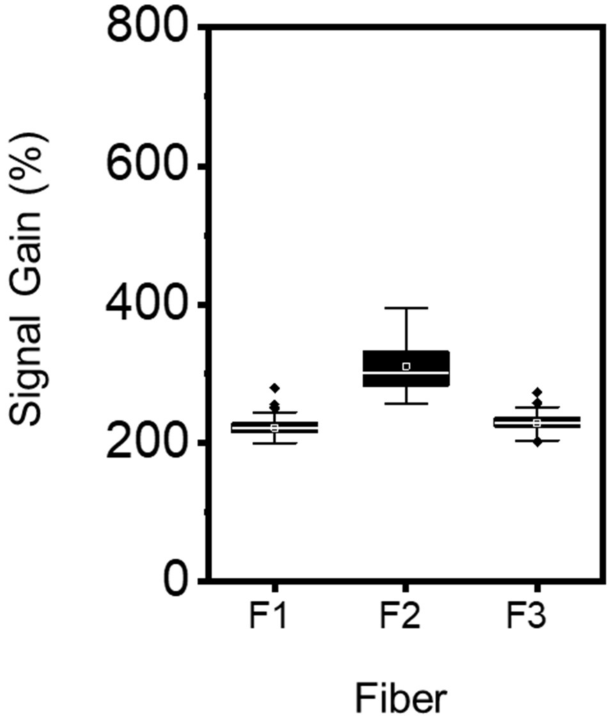

**Supplementary Figure S13:** Fiber-to-fiber signal gain variance in the presence of 8 μM dopamine (CV = 0.142). Error bars represent the standard deviation.

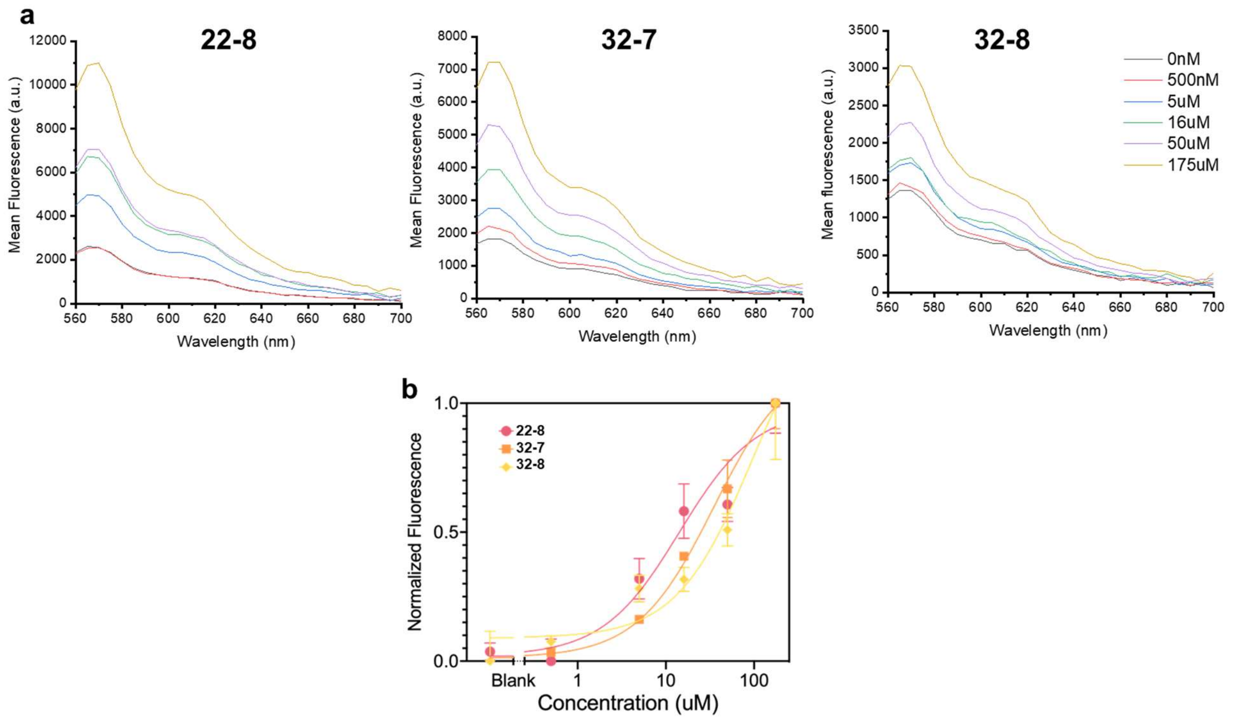

**Supplementary Figure S14: Raw signal analysis of selected cortisol DBS constructs.** Representative concentration-dependent emission spectra of various cortisol DBS constructs. Bottom panel shows fluorescence at peak emission (565 nm). Plots show raw fluorescence as a function of cortisol concentration. All plots show averages from three replicates. Error bars represent the standard deviation.

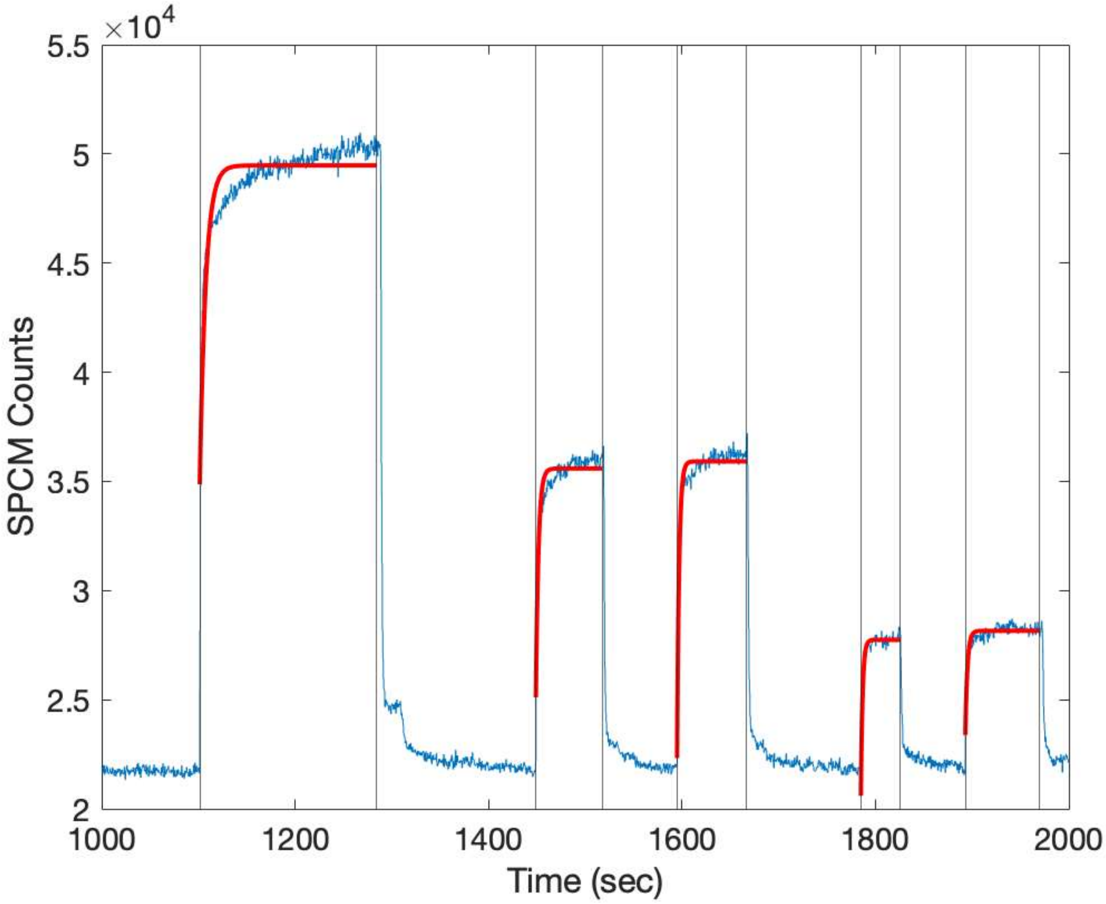

**Supplementary Figure S15: Kinetics fits from cortisol real-time measurements.** Fittings failed to converge to a single exponential due to fast on/off switching rates of the aptamer. These fits estimate $k_{obs}$ to be in the range of 0.1-0.6 $s^{-1}$, which cannot be confidently resolved with the 500 ms integration time of our system.

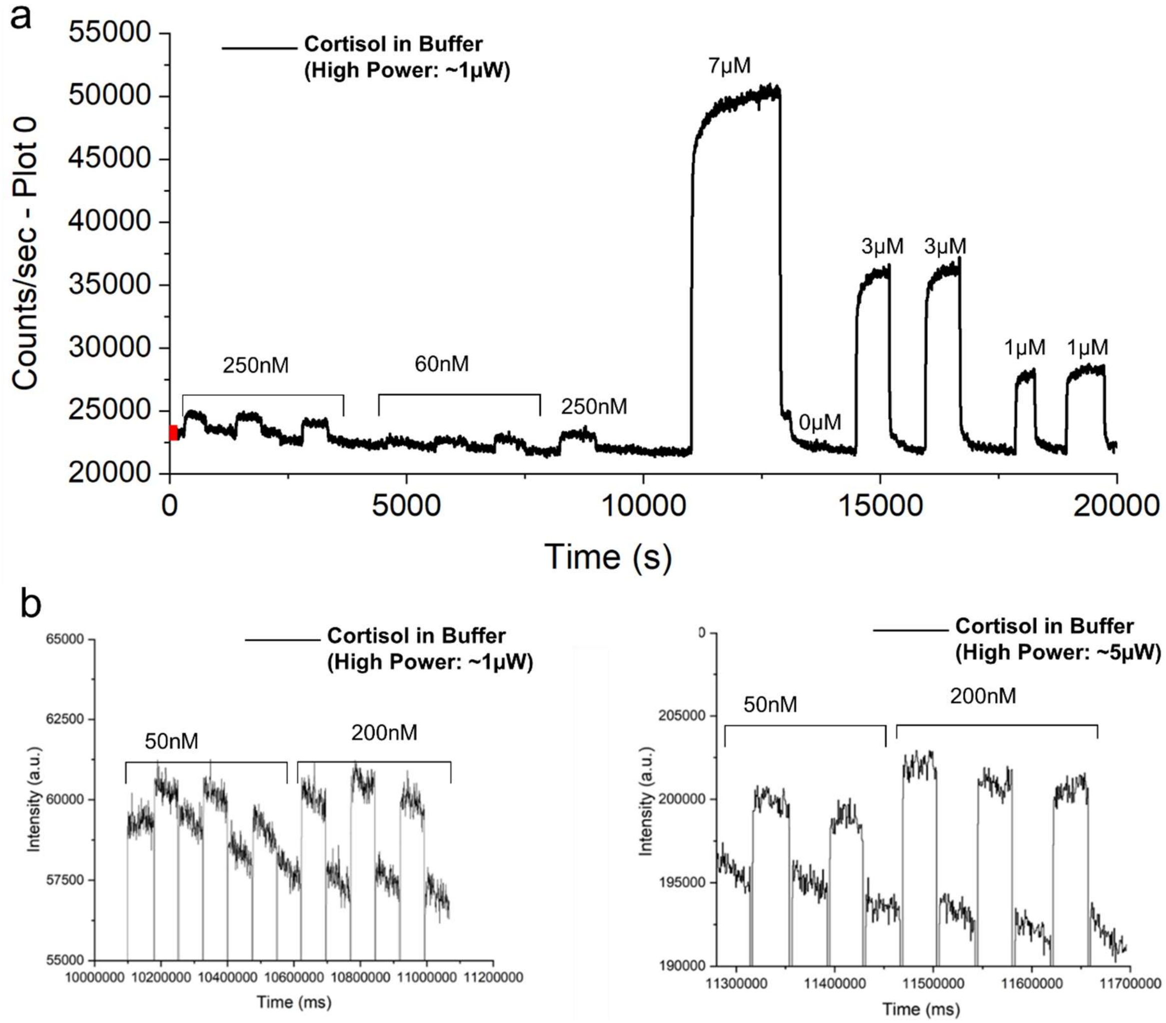

**Supplementary Figure S16: Additional raw data showing real-time measurements of cortisol** at different concentrations in buffer at ~1 μW and ~5 μW laser power.

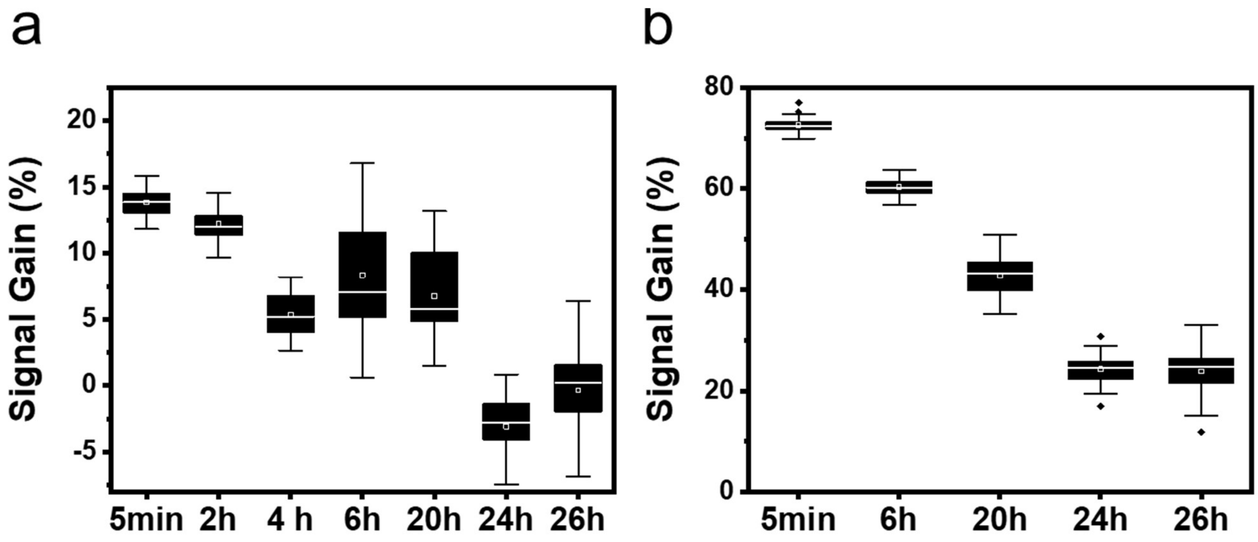

**Supplementary Figure S17: Signal enhancement after undiluted plasma exposure at different time points.** Probes were exposed to undiluted human plasma for different intervals of time (5 min, 2 h, 4 h, 6 h, 20 h, 24 h and 26 h), and then transferred to either **a**, plasma or **b**, buffer containing 5 µM cortisol. Error bars represent the standard deviation.

**Supplementary Table 1:** Sequences used in this work. Black text represents the $L_{Bubble}$ linker, bold text represents the anchor strand, orange text represents the aptamer, underlined text represents a mismatch, and red text represents displacement strand with IowaBlack-FQ quencher at the 5' end.

| | | | |
|---|---|---|---|
| **Dopamine aptamer strand** | | | **AAATACAACAAGAAAAAACTCTCGGGACGACG/i-Cy3/CCAGTTTGAAGGTTCGTTCGCAGGTGTGGAGTGACGTCGTCCC** |
| **Dopamine aptamer strand with mismatch** | | | **AAATACAACAAGAAAAAACTCTCGGGACGGCG/i-Cy3/CCAGTTTGAAGGTTCGTTCGCAGGTGTGGAGTGACGTCGTCCC** |
| Bubble | **DS** | **Anchor** | **Sequence (5' → 3')** |
| 22 | 7 | 18 | GTCGTCCTCTTCACATATCTA**TTTTTTCTTGTTGTATTT** |
| 22 | 8 | 18 | CGTCGTCCTCTTCACATATCTA**TTTTTTCTTGTTGTATTT** |
| 22 | 9 | 18 | CGTCGTCCCCTCTTCACATATCTA**TTTTTTCTTGTTGTATTT** |
| 22 | 10 | 18 | CGTCGTCCCGCTCTTCACAATATCTA**TTTTTTCTTGTTGTATTT** |
| 22 | 11 | 18 | CGTCGTCCCGATTCTTCCACAATATCTA**TTTTTTCTTGTTGTATTT** |
| 22 | 12 | 18 | CGTCGTCCCGAGCTCTTTCCACAATATCTA**TTTTTTCTTGTTGTATTT** |
| 32 | 8 | 18 | CGTCGTCCTCTCCACTATACTCATAATATCTA**TTTTTTCTTGTTGTATTT** |
| 32 | 9 | 18 | CGTCGTCCCTCTTCCACTATACTCATAATATCTA**TTTTTTCTTGTTGTATTT** |
| 32 | 10 | 18 | CGTCGTCCCGCTCTTCCACTATACTCATAATATCTA**TTTTTTCTTGTTGTATTT** |
| 32 | 11 | 18 | CGTCGTCCCGATTCTTCCACTATACTCATAAATATCTA**TTTTTTCTTGTTGTATTT** |
| 32 | 12 | 18 | CGTCGTCCCGAGCTCTTTCCACTATACTCATAAATATCTA**TTTTTTCTTGTTGTATTT** |
| 12 | 8 | 18 | CGTCGTCCTCTA**TTTTTTCTTGTTGTATTT** |
| 12 | 9 | 18 | CGTCGTCCCTTCTA**TTTTTTCTTGTTGTATTT** |
| 12 | 10 | 18 | CGTCGTCCCGTCTCTA**TTTTTTCTTGTTGTATTT** |
| 12 | 11 | 18 | CGTCGTCCCGACTCTCTA**TTTTTTCTTGTTGTATTT** |
| 12 | 12 | 18 | CGTCGTCCCGAGCTCTTCTA**TTTTTTCTTGTTGTATTT** |
| **Cortisol aptamer strand** | | | **AGAAAATACAACAAGAAAAAACTCTCGGGACGAC/iCy3/GCCAG AAGTTTACGAGGATATGGTAACATAGTCGTCCC** |
| Bubble | **DS** | **Anchor** | **Sequence (5' → 3')** |
| 22 | 8 | 18 | TCCCGAGAGAACATCCTCACACACCCA**TTTTCTTGTTGTATTTTCT** |
| 32 | 7 | 18 | CCCGAGAGAACATCCTCACCTAACACATTACACCCA**TTTTCTTGTTGTATTTTCT** |
| 32 | 8 | 18 | TCCCGAGAGAACATCCTCACCTAACACATTACACCCA**TTTTCTTGTTGTATTTTCT** |

**Supplementary Table 2: Effect of $L_{Bubble}$ and $L_{DS}$ on effective binding affinity for dopamine.** (n = 3 replicates).

| Construct | $K_D^{eff}$ ( $\mu M$) | $Y_{min}$ (RFU) | $B_{max}$ (RFU) | $R^2$ |
|---|---|---|---|---|
| **12-8** | 135.5 | 2067 | 11,227 | 0.9866 |
| **12-9** | 103.4 | 1,457 | 3,188 | 0.4053 |
| **12-10** | 177.1 | 1,297 | 3,277 | 0.9453 |
| **12-11** | 82.4 | 1,751 | 10,356 | 0.9711 |
| **12-12** | 103.8 | 644.2 | 1,604 | 0.9811 |
| **22-7** | 36.5 | 2,180 | 7,220 | 0.984 |
| **22-8** | 58.73 | 11,919 | 16,856 | 0.9173 |
| **22-9** | 72.0 | 4,240 | 16,328 | 0.9882 |
| **22-10** | 68.31 | 2,824 | 13,938 | 0.9797 |
| **22-11** | 113.9 | 1,419 | 11,179 | 0.9824 |
| **22-12** | 169.0 | 1,042 | 4,699 | 0.9637 |
| **32-8** | 114.3 | 10,067 | 15,042 | 0.9446 |
| **32-9** | 66.22 | 6,031 | 16,951 | 0.9862 |
| **32-10** | 48.89 | 5,593 | 17,603 | 0.9674 |
| **32-11** | 104.2 | 2,351 | 16,481 | 0.9865 |
| **32-12** | 146.4 | 1,146 | 10,916 | 0.932 |

**Supplementary Table 3: Effect of $L_{Bubble}$ and $L_{DS}$ lengths on kinetics ($\tau_{obs}$) for dopamine.** (n = 3 replicates).

| Construct | $\tau_{obs}$(sec) |
|---|---|
| **12-8** | 173.1 |
| **12-11MM** | 35.22 |
| **12-11** | 445.8 |
| **22-7** | 2.95 |
| **22-8** | 86.27 |
| **22-8 MM** | 121.1 |
| **22-9** | 65.39 |
| **22-10** | 171.7 |
| **22-11** | 1032 |
| **32-8** | 226.5 |
| **32-9** | 169.5 |
| **32-10** | 354.2 |
| **32-11** | 927.4 |

**Supplementary Table 4:** Signal gain from dopamine fiber measurements in buffer

| Concentration ($\mu$M) | N (Replicates) | Mean Signal Gain (%) | CV |
|---|---|---|---|
| 1 | 5 | 28.9 | 0.036 |
| 8 | 16 | 195 | 0.229 |
| 80 | 5 | 671 | 0.118 |
| 200 | 2 | 1,310 | 0.202 |
| 800 | 2 | 2,056 | 0.161 |

**Supplementary Table 5:** Kinetics fits for dopamine fiber measurements in buffer

| Concentration | N | Mean $k_{obs}$ ($sec^{-1}$) | CV |
|---|---|---|---|
| 8 | 4 | 0.094733 | 0.167718 |
| 80 | 3 | 0.117986 | 0.244692 |
| 200 | 2 | 0.141867 | 0.330711 |
| 800 | 2 | 0.208099 | 0.029817 |

**Supplementary Table 6:** Signal gain from dopamine and cortisol fiber measurements in aCSF or plasma

| Concentration ($\mu$M) | N (Replicates) | Mean Signal Gain (%) | CV |
|---|---|---|---|
| **Dopamine (aCSF)** | | | |
| 0.5 | 3 | 6.7 | 0.005 |
| 1 | 6 | 11.8 | 0.033 |
| 8 | 3 | 141 | 0.032 |
| 80 | 4 | 771 | 0.093 |
| 200 | 1 | 1,228 | n/a |
| 800 | 1 | 1,527 | n/a |
| **Cortisol (Plasma)** | | | |
| 0.2 | 3 | 6.5 | 0.014 |
| 1.4 | 3 | 18.2 | 0.003 |
| 10 | 5 | 57.6 | 0.188 |
| 140 | 3 | 296 | 0.033 |

**Supplementary Table 7: Effect of $L_{Bubble}$ and $L_{DS}$ on effective binding affinity for cortisol.** (n = 3 replicates).

| Construct | $K_D^{eff}$ ( $\mu M$) | $Y_{min}$ (RFU) | $B_{max}$ (RFU) | $R^2$ |
|---|---|---|---|---|
| **22-8** | 14.24 | 2655 | 10034 | 0.9059 |
| **32-8** | 97.46 | 1496 | 3644 | 0.8930 |
| **32-7** | 35.03 | 2011 | 8071 | 0.9780 |

**Supplementary Note 1**

In order to predict the behavior of our DBS constructs, we assume that a population of constructs exists in equilibrium between quenched (Q), unfolded (U), and target-bound (B) states, similar to the analysis in (similar to the analysis in Wilson *et al.* [16]). In the Q state, two double-stranded regions form between the aptamer and displacement probes, bringing the quencher and fluorophore in close proximity, yielding an OFF state that produces no fluorescent signal. For both the U and B states, the aptamer is single-stranded, with the quencher farther from the fluorophore and thus yielding a signal-ON state.

In the absence of target, the DBS construct is in equilibrium between Q and U states, with the U constructs contributing to background signal. This equilibrium is dictated by the kinetics of the double-stranded displacement region hybridizing ($k_{on}^{DS}$) and unhybridizing ($k_{off}^{DS}$), as described by Equation S1.

$$[S1]\ K_Q = \frac{[Q]}{[U]} = \frac{k_{on}^{DS}}{k_{off}^{DS}}$$

In the presence of target, the DBS establishes an equilibrium between Q, U, and B states. The equilibrium between U and B states is dictated by the kinetics of the native aptamer binding ($k_{on}^{apt}$) and unbinding ($k_{off}^{apt}$) as described by Equation S2.

$$[S2]\ K_D^{apt} = \frac{[U][T]}{[B]} = \frac{k_{off}^{apt}}{k_{on}^{apt}}$$

The total expected signal (S) at a given target concentration is proportional to the aptamer concentration [apt] and the sum of the unfolded ($F_U$) and target-bound ($F_B$) fractions of aptamer as described by Equation S3.

$$[S3]\ S = [apt] * (F_U + F_B) = [apt] * \left(\frac{[U]}{[Q]+[U]+[B]} + \frac{[B]}{[Q]+[U]+[B]}\right)$$

Combining equations S1 – S3, we can derive expected signal as a function of target concentration, $K_Q$, and $K_D^{apt}$.

$$S = [apt] * \left(\frac{[U]+[B]}{[Q]+[U]+[B]}\right)$$

$$S = [apt] * \frac{K_D^{apt} + [T]}{K_D^{apt}(1+K_Q) + [T]}$$

$$[S4]\ S = [apt] * \frac{[T]}{K_D^{apt}(1+K_Q) + [T]} + \frac{[apt]}{(1+K_Q)}$$

The first term in Equation S4 describes the target-dependent signal, while the second term describes background. From this derivation, it becomes clear that background is inversely proportional to ($1+K_Q$) and increasing $K_Q$ will decrease background. $K_Q$ is expected to increase with increasing displacement strand length ($L_{DS}$) as it becomes more energetically favorable to form a double-stranded region or decreasing $L_{bubble}$ length as the effective concentration of the displacement strand is increased. We can define the effective dissociation constant ($K_D^{eff}$) of the construct as a function of $K_Q$ and $K_D^{apt}$ described by equation S5.

$$[S5]\ K_D^{eff} = K_D^{apt}(1 + K_Q)$$

We assume that $K_D^{apt}$ is fixed as a function of the native aptamer; however, increasing $K_Q$ can increase the $K_D^{eff}$ and shift our binding curve to the right.

Additionally, we can modify the overall kinetics of our construct by tuning $L_{DS}$ or $L_{bubble}$. It has been shown that $k_{on}^{DS}$ scales inversely with $L_{bubble}$, while changing $L_{bubble}$ has little effect on $k_{off}^{DS}$. Additionally, decreasing $L_{DS}$ will decreasing hybridization strength of the double stranded region and can be predicted to increase $k_{off}^{DS}$ more than $k_{on}^{DS}$, Therefore, we can expect to increase the overall $k_{obs}$ from the Q to the B state by decreasing $L_{bubble}$ or decreasing $L_{DS}$. By testing several $L_{bubble}$ and $L_{DS}$ values, we were able to choose an optimal construct with the desired dynamic range and response time.
